# Contemporary hybridization and localized genomic differentiation between grey seal subspecies

**DOI:** 10.64898/2026.08.10.743471

**Authors:** Anastasia Konstantopoulou, Ari Löytynoja, Theresa Koller, Emmi Olkkonen, Anders Galatius, Morgan L. McCarthy, Iben Stokholm, Sandra M. Granquist, Bjørn Munro Jenssen, Mart Jüssi, Ivar Jüssi, Mervi Kunnasranta, Ursula Siebert, Ailsa Hall, Petri Auvinen, Jukka Jernvall, Rune Dietz, Jonas Teilmann, Claudius F. Kratochwil, Morten Tange Olsen

## Abstract

Grey seals are divided into two subspecies, *Halichoerus grypus grypus* in the Baltic Sea and *H. grypus atlantica* in the North Atlantic Ocean. Historically, intense hunting caused local extinctions of grey seals across their range and promoted geographical isolation of the subspecies. However, recent population recovery and recolonization have renewed their overlap in a contact zone in Danish and Swedish waters. Here, using whole-genome sequencing data from 119 individuals, we investigate the demographic history of grey seals, the divergence of the subspecies, and genomic signatures of local adaptation and potential hybridization in the contact zone. We estimate that divergence began approximately 10,000 years ago, with gene flow ceasing 2000 years ago. We detect peaks of high genetic differentiation near genes with putative functions in osmo- and thermoregulation, consistent with salinity and temperature differences between the Baltic Sea and the North Atlantic. As evidence of the geographic isolation breaking up, we report a hybrid individual in the southwest Baltic contact zone, with Baltic maternal and Atlantic paternal ancestry, and identify a migrant of Baltic origin in the North Sea. Our study illustrates how local environmental variation and long-term hunting pressure have contributed to divergence between mammal subspecies over a short evolutionary timescale, with recent population recovery, recolonization and hybridization reshaping gene flow dynamics. Broadly, our work emphasizes the importance of studying evolutionary processes amid anthropogenic influences, including both disturbances and conservation successes, where demographic fluctuations, range shifts, altered gene flow, and adaptation to changing environments interact in complex, potentially consequential ways.

## 1. Introduction

Hybridization is a multifaceted evolutionary process (Abbott et al. 2016; Runemark et al. 2019), with outcomes including the merging of the hybridizing populations (e.g., Taylor et al., 2006), introgression of potentially adaptive alleles (e.g., Valencia-Montoya et al., 2020), formation of stable hybrid zones (e.g., Poelstra et al., 2014), as well as the emergence of new species (Abbott et al. 2013; e.g., Meier et al. 2017; Lopes et al. 2023). Contemporary hybridization events allow us to observe this process in its early stages, before its long-term evolutionary outcomes unfold. Studying hybridization in natural systems can help detect genes under selection and explore the genetic basis of reproductive isolation, identify genomic regions derived from introgression, and inform decisions in conservation (Abbott et al. 2016). In pinnipeds and other marine mammals, hybridization has been frequently reported (Kovacs et al. 1997; Bérubé and Aguilar 1998; Lancaster et al. 2006; Schaurich et al. 2012; Savriama et al. 2018; Lopes et al. 2021; Lopes et al. 2023), possibly reflecting their high dispersal ability and weak prezygotic isolation (Lopes et al. 2023).

The grey seal (*Halichoerus grypus* Fabricius, 1791) has a cold temperate to sub-Arctic distribution and includes three main genetic populations: in the North-west (NW) Atlantic, the Northeast (NE) Atlantic and the Baltic Sea (Boskovic et al. 1996; Klimova et al. 2014; McCarthy et al. 2025). Two subspecies of grey seals are currently recognized by the IUCN and the Society for Marine Mammalogy: the Atlantic grey seal (*H. g. atlantica*), which consists of the NW and NE Atlantic populations, and the Baltic grey seal (*H. g. grypus*) (Berta and Churchill 2012; Olsen et al. 2016). The divergence between Baltic and NE Atlantic grey seals has been linked to the formation of the Baltic Sea following the Last Glacial Maximum (Klimova et al. 2014; Fietz et al. 2016), yet its timing and underlying drivers have not been assessed using whole-genome sequencing (WGS) data.

The modern Baltic Sea environment is considered the world’s largest brackish-water estuary and likely developed into its present form only after complete deglaciation, with the Littorina Sea phase beginning around 8500–8000 years ago (Björck 1995; Andrén et al. 2011). This created the low-salinity conditions [15-18 psu (practical salinity units) at the entrance to the North Sea, 0-2 psu in the innermost parts; HELCOM 2023] that, together with the occurrence of seasonal sea-ice and low winter temperatures [minimum annual SST (satellite sea surface temperature) of -0.3 to 3 °C in the Baltic and 8 °C near the North Sea entrance to the Baltic; Stramska and Białogrodzka 2015], distinguish its marine ecosystem and habitats from those of the neighboring NE Atlantic. Such environmental differences have been suggested to contribute to local genetic adaptation in many Baltic Sea marine organisms, including marine mammals (e.g., Johannesson and André 2006; Autenrieth et al. 2024; Celemín et al. 2025). The Baltic subspecies, unlike the Atlantic, mainly breeds on sea-ice, and Baltic and NE Atlantic grey seals have asynchronous pupping seasons, which can be at least partially attributed to plasticity in breeding phenology (Härkönen et al. 2007; Bowen et al. 2020; Galatius et al. 2024). While overall genetic differentiation between the subspecies is low (FST=0.034-0.057; McCarthy et al. 2025), it is possible that they exhibit localized genomic regions of high differentiation that also contribute to the observed differences in their reproductive biology and are potentially associated with adaptation to local environmental conditions (e.g., osmo- or thermoregulation).

Across the species’ range, grey seal populations suffered severe declines due to hunting and other anthropogenic pressures. This led to loss of genetic lineages as early as in the Mesolithic (Ahlgren et al. 2022) and local population extinctions along the European coastlines from the Middle Ages to the 19th century (Härkönen et al. 2007; Olsen et al. 2018). Specifically, hunting pressure in the contact region of the two subspecies may have reduced gene flow and reinforced their separation (Fietz et al. 2016). Upon gradual protection in the mid-20^th^ century, the species has shown a rapid recovery (Härkönen et al. 2007; Galatius et al. 2024; Vanko et al. 2026) and has started recolonizing areas of its historical range (Härkönen et al. 2007; Fietz et al. 2016; Galatius et al. 2020; Galatius et al. 2024). As a result, the distributions of the Baltic and Atlantic grey seals are starting to overlap, with intermediate Danish and Swedish waters constituting a contact zone (Fietz et al. 2016; Galatius et al. 2024). This raises the question of whether hybridization among the two subspecies is common, and if it affects gene flow dynamics through the occurrence of backcrosses. Generally, it can be assumed that the frequency of hybridization and its effect on gene flow will be largely influenced by the fitness of first-generation hybrids. Hybrid fitness may be, for example, affected by genetic incompatibilities, as well as by alleles involved in local adaptation to the contrasting environments experienced by the Baltic and Atlantic populations (Harrison and Larson 2016). Previous studies, using limited datasets of mtDNA and microsatellite markers (Fietz et al. 2016) as well as ddRADseq data (McCarthy et al. 2025), hinted at shifts in subspecies boundaries and possible hybridization at haul-out sites in Southwest (SW) Baltic. However, as whole-genome information has been lacking so far, supporting evidence is limited.

Here, we investigate the long-term demographic history of grey seals, the divergence of the NE Atlantic and Baltic populations, their contemporary hybridization, and localized genomic regions of high differentiation that may reflect adaptive divergence. To this end, we analyze WGS data from 119 individuals, including data from pups from the Rødsand haul-out site in the Danish part of the SW Baltic that constitutes a likely contact zone of the two subspecies (Figure 1A; Table 1). Moreover, possibly admixed individuals have been previously reported at this site (Fietz et al. 2016; McCarthy et al. 2025). After inferring the demographic history and timing of divergence of the populations, we explore the subspecies’ current population structure and assess hybridization. Next, we investigate the genomic composition of the identified hybrid individual, including its maternal and paternal origins. Finally, we characterize genomic regions of high differentiation between the subspecies and evaluate whether the functions of nearby genes are consistent with local adaptation to contrasting salinity and temperature conditions.

**Table 1.** Sample information summary. Individuals used in the analyses are included. For each sampling location, indicated by its site code, the number of samples (including the total number per group) and their subspecies classification is provided. The ringed seal samples from Greenland are also listed.

| Species | Subspecies | Sampling Location | Site code | Number of samples |
| --- | --- | --- | --- | --- |
| Grey Seal | Atlantic | Denmark, Thyborøn | DEN_THY | 17 |
| Grey Seal | Atlantic | Iceland, Breiðafjörður | ICE_BRE | 4 |
| Grey Seal | Atlantic | Iceland, Patreksfjörður | ICE_PAT | 1 |
| Grey Seal | Atlantic | Iceland, Strandir | ICE_STR | 4 |
| Grey Seal | Atlantic | Iceland, Djúpivogur | ICE_DJU | 7 |
| Grey Seal | Atlantic | Norway, Froan | NOR_FRO | 9 |
| Grey Seal | Atlantic | Russia, Kola | RUS_KOL | 9 |
| Grey Seal | Atlantic | UK, Isle of May | UK_ILM | 5 |
| Grey Seal | Atlantic |  |  | 56 |
| Grey Seal | Baltic | Denmark, Christiansø | DEN_CHR | 20 |
| Grey Seal | Baltic | Denmark, Rødsand | DEN_ROD | 20 |
| Grey Seal | Baltic | Estonia, Gulf of Riga | EST_GOR | 5 |
| Grey Seal | Baltic | Finland, Bothnian Bay | FIN_BOB | 4 |
| Grey Seal | Baltic | Finland, Gulf of Finland | FIN_GOF | 4 |
| Grey Seal | Baltic | Sweden, Falsterbo | SWE_FAL | 5 |
| Grey Seal | Baltic | Sweden, Stockholm Archipelago | SWE_STA | 5 |
| Grey Seal | Baltic |  |  | 63 |
| Grey Seal | All |  |  | 119 |
| Ringed Seal | Arctic | Greenland, Itoqqortoormiit | GRE_IQ | 7 |
| Ringed Seal | Arctic | Greenland, Qaanaaq | GRE_QA | 10 |
| Ringed Seal | Arctic |  |  | 17 |

**Figure 1.**
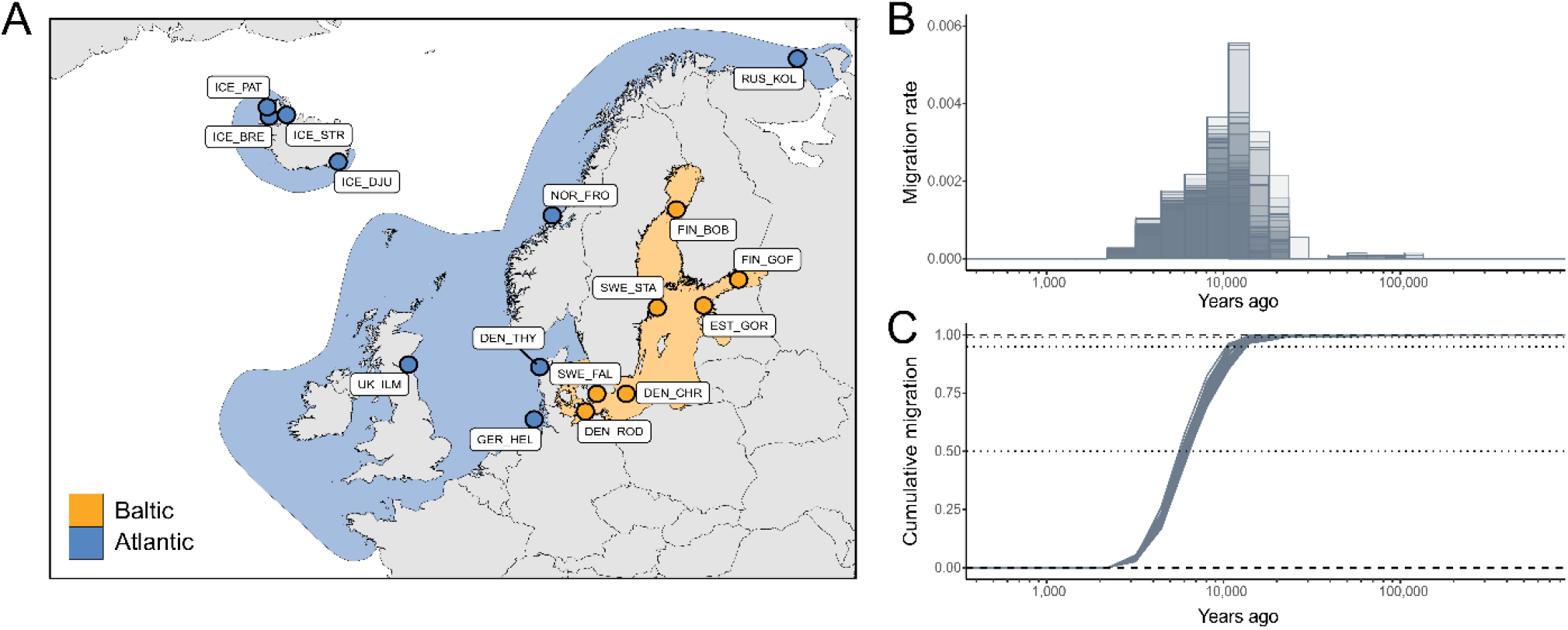
Sampling localities and the history of subspecies divergence. A) Sampling locations (points) and distributions of the Baltic and NE Atlantic populations (shading). B) Estimated migration rate (m) between pairs of individuals from the Baltic and NE Atlantic populations through time. C) Estimated cumulative migration (M) between pairs from the same populations. (B) and (C) indicate a decline in gene flow beginning approximately 10,000 years ago, a cessation around 2000 years ago and an estimated population split time of ∼6000 years ago (M=50%).

## 2. Results

### 2.1 Demographic history and divergence time

We inferred demographic histories and the split time of the NE Atlantic and Baltic grey seals and tested whether it correlates with the postglacial history of the Baltic Sea by estimating coalescence rates, converting them to estimates of effective population size (*N*_e_) and fitting a continuous isolation-with-migration model. The *N*_e_ of grey seals started declining around 150,000 years ago, broadly coinciding with the later part of the penultimate glacial period. *N*_e_ dropped below 5000 around 50,000 years ago (Figure S1), during the middle Weichselian glaciation. Interestingly, although low, the inferred *N*_e_ of grey seals did not drop below 2500 at any point in the species’ history, suggesting long-term persistence through Late Pleistocene glacial cycles. The *N*_e_ trajectories of NE Atlantic and Baltic grey seals separated around 10,000 years ago, broadly coinciding with the final retreat of the Weichselian ice sheet and the early postglacial development of the Baltic basin. The NE Atlantic population showed a slight increase in *N*_e_, whereas the Baltic population remained small. The migration rate analysis also revealed a drop in gene flow around that time (Figure 1B). The cumulative migration rate (M), using the cutoff of 50%, suggested a median population split time of 6000 years ago (Figure 1C), while gene flow completely ceased around 2000 years ago (Figure 1B). These estimates broadly coincide with the establishment of the brackish Littorina Sea during the Mid Holocene (Andrén et al. 2011) and the Iron Age, respectively, when human exploitation of grey seals was already prevalent (Ahlgren et al. 2022). Notably, an increase in *N*_e_ was observed in the last 1500 years for both subspecies (Figure S1); however, estimates for timescales as recent as few thousand years ago are commonly not reliable and should be interpreted with caution (Hilgers et al. 2025).

In summary, our WGS-based demographic analyses indicate a gradual divergence between Baltic and NE Atlantic grey seals, characterized by progressive reductions in gene flow over time. This pattern broadly overlaps with the postglacial emergence of distinct Baltic environmental conditions, and possibly early human influence during the Stone and Iron Age.

### 2.2 Genetic differentiation and population structure

We studied the population structure of the grey seal subspecies using Principal Component Analysis (PCA), Identity-by-state (IBS) distance estimation and admixture analysis. The PCA separated the Baltic from the NE Atlantic subspecies (Figure 2A), with the Atlantic individuals further clustering according to their geographic origin along PC2. One individual from the Rødsand haul-out site, HG301, clustered in the middle of the two groups, consistent with a hybrid origin. HG517, a pup sampled at Thyborøn, Denmark, clustered within the Baltic subspecies, suggesting migration from the Baltic into the North Sea. The admixture analysis and IBS distances further confirmed the PCA results (Figure 2B; Figure S2). Interestingly, grey seals sampled at the Rødsand site (DEN_ROD) appeared to form a distinct cluster, being highly genetically similar to each other, as supported by the PCA, admixture analysis, and very low IBS distances between pairs from this population (Figure S2). KING-based kinship estimates supported elevated relatedness in this site (Table S1), consistent with increased rates of inbreeding.

**Figure 2.**
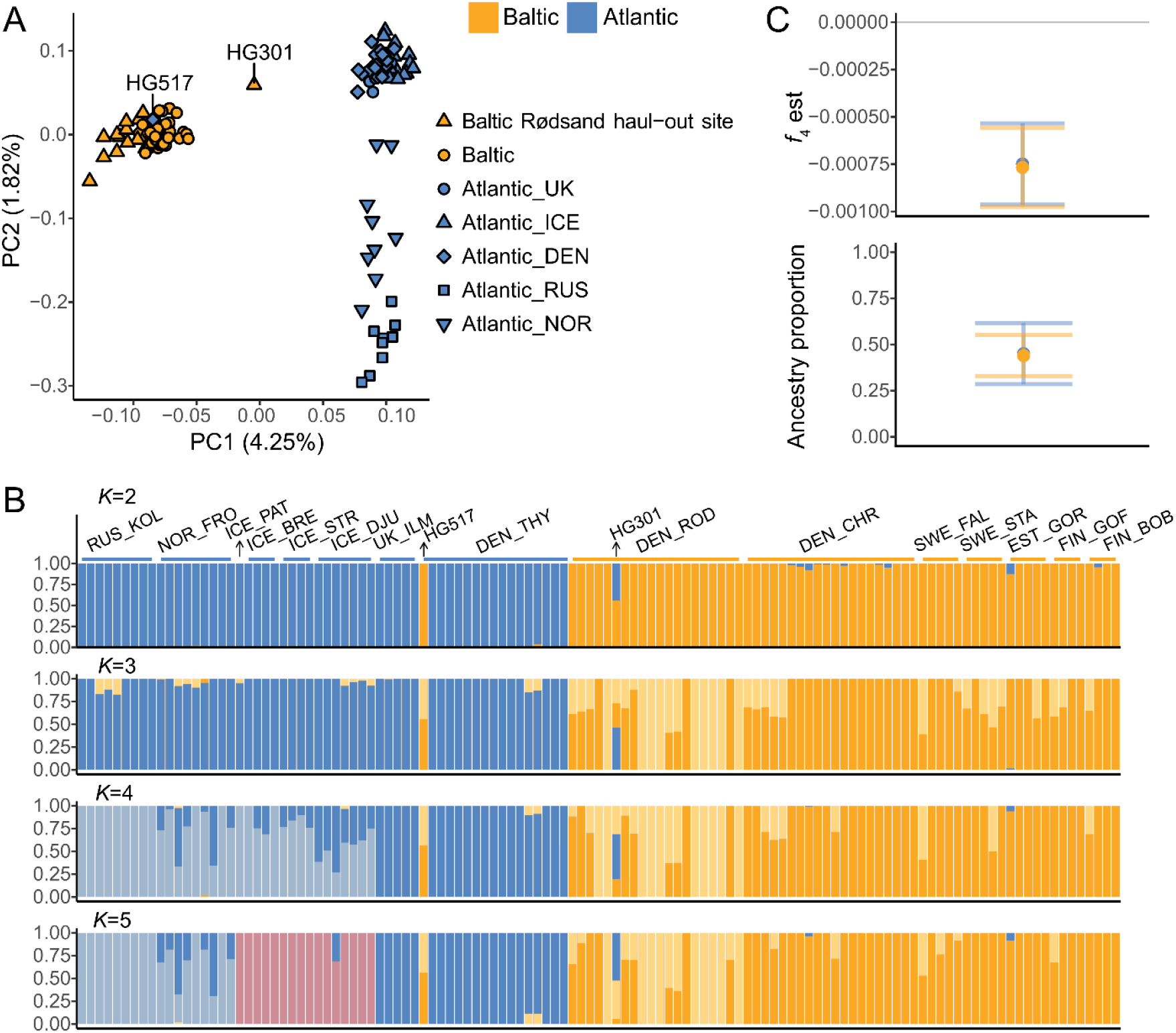
Population structure and hybrid identification. A) PCA plot showing the clustering of individuals along the first two PCs, separating NE Atlantic and Baltic grey seals and revealing a hybrid (HG301) and a Baltic-like migrant individual (HG517). B) Admixture analysis for *K*=2-5 genetic clusters, congruent with the PCA results. Names on top of the plot correspond to codes of sampling sites (Table 1), each bar represents one individual, and colors denote different ancestries. HG301 and HG517 are marked. C) Top panel: estimates of *f*_4_-statistics supporting introgression into HG301 from both the Baltic and the Atlantic. Bottom panel: ancestry proportions of HG301 from the Baltic and the Atlantic as estimated by the *f*_4_-ratio test (mean±SE), with both proportions close to 50%.

Overall, the PCA, admixture analysis, and IBS distances consistently separated Baltic and NE Atlantic grey seals, while also revealing geographic structure within the Atlantic subspecies. These analyses further provide evidence for ongoing hybridization, migration, and distinct clustering of Baltic grey seals from the Danish Rødsand population.

### 2.3 Validation of hybrid ancestry and parental origins

Building on the identification of HG301 as a putative hybrid individual, we formally tested its hybrid origin and ancestry proportions using *f*_4_-statistics. The *f*_4_-statistic was significantly negative (Figure 2C; *P*<0.001) when testing for gene flow from both the Baltic and the Atlantic, indicating admixture from both subspecies into HG301. In the *f*_4_-ratio test, the estimates of ancestry proportion (α) were near 0.5 and significant (Figure 2C; Figure S3; Baltic: α=0.448, SE=0.111, z>3; Atlantic: α=0.457, SE=0.14, z>2). These roughly equal ancestry proportions are compatible with expectations for a first-generation hybrid.

Next, we analyzed sex-linked variation to infer parental contributions in the hybrid. To achieve this, molecular sex was first assigned to individual seals by identifying sex-linked contigs. The final set of X contigs had a total length of 117.492 Mbp, approximating that of the dog X chromosome (∼127 Mbp; ROS_Cfam_1.0, Ensembl release 115). Sexes could be clearly separated based on the mean coverage of the X-linked contig *H. fasciata* 209 (Hfa209) (Figure 3A). The hybrid individual, HG301, was inferred to be a male and the migrant, HG517, was determined to be a female. In a PCA including only X-linked contigs and only male individuals, the hybrid’s X contigs clustered with the Baltic (Figure 3B), suggesting that the maternal X chromosome likely originated from the Baltic and, thus, that the paternal population was NE Atlantic.

**Figure 3.**
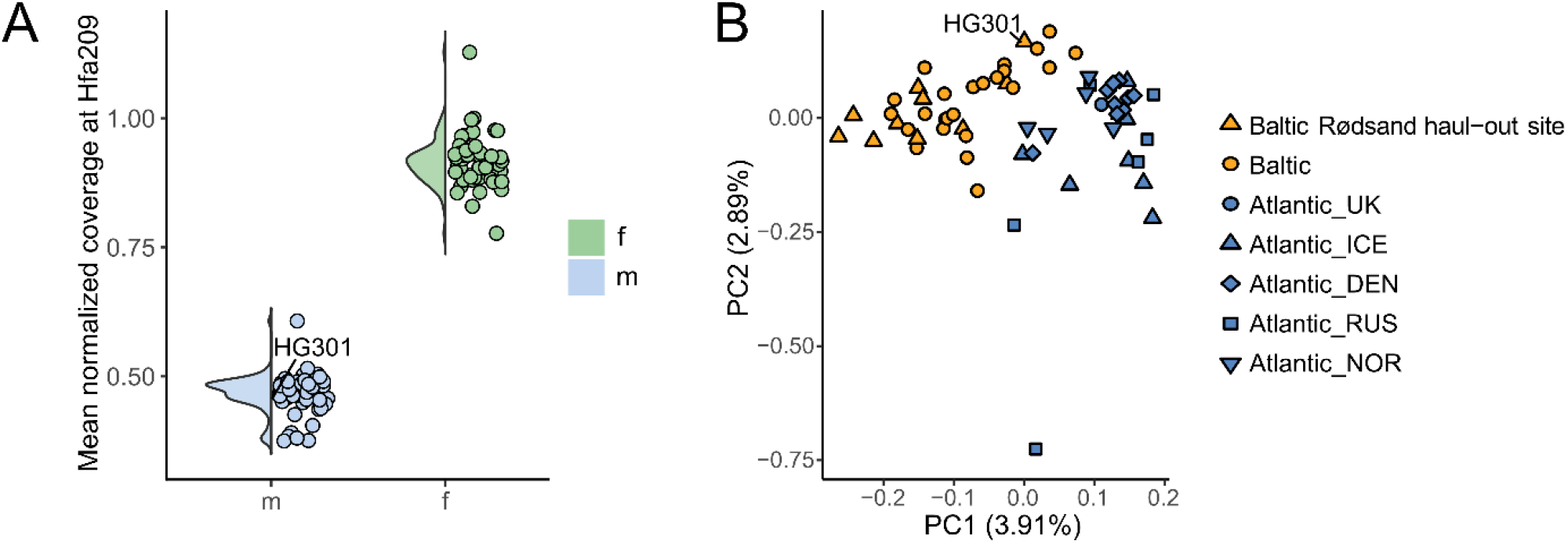
Sex assignment and inference of hybrid’s maternal population. A) Sex identification based on the mean coverage of windows along the X-linked contig Hfa209, normalized by the mean coverage of the five largest autosomal contigs. Males, including the hybrid HG301, show around half the female coverage. B) PCA analysis on the X-linked contigs of a male-only data set. HG301 clusters near Baltic males, suggesting that its maternal X chromosome (mother) was Baltic.

In summary, the *f*_*4*_-statistic and *f*_*4*_-ratio test confirmed that HG301 has mixed Baltic and NE Atlantic ancestry, with approximately equal contributions from both subspecies, consistent with a first-generation hybrid. Sex-linked analyses inferred HG301 to be male and suggested a Baltic maternal ancestry, implying a NE Atlantic paternal contribution.

### 2.4 Genomic signals of differentiation and potential local adaptation

We assessed potential local adaptation in the two subspecies by screening for localized genomic regions marked by high allele frequency differences (ΔAF peaks). The overall genetic differentiation between the populations was low: the ΔAF distribution was skewed towards lower values, with a mean value of 0.028 and a median of 0.001. While no alleles are fixed between subspecies, the 99.99^th^ percentile was 0.51 and the maximum ΔAF reached 0.74, suggesting that some sites exhibit exceptionally elevated levels of differentiation. The highest ΔAF was observed on contigs Hfa027 and Hfa020 (dog chromosomes 7 and 8; Figure 4A,B,C). We used ΔAF≥0.55 as a cutoff to capture the regions of greatest differentiation, focusing on windows of high ΔAF rather than on isolated variants (Figure 4A). The selected set corresponded to the top ∼0.004% of the ΔAF distribution and comprised 459 SNPs, indicating highly localized genomic differentiation between the subspecies despite low genome-wide divergence.

**Figure 4.**
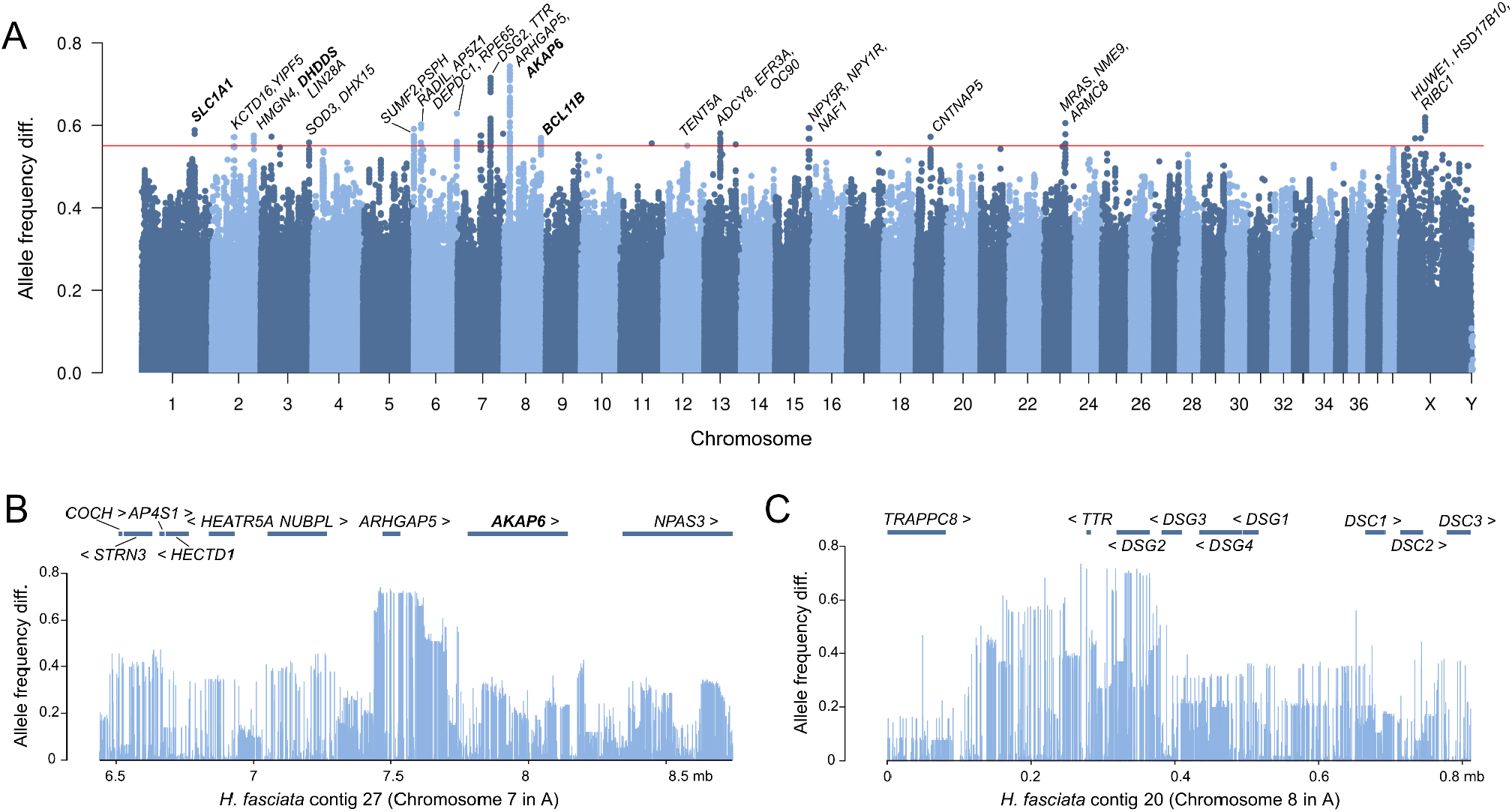
Candidate regions of differential adaptation between the subspecies. A) Allele frequency differences (ΔAF) between individuals grouped by subspecies (Baltic and Atlantic) at sites that were successfully lifted over to a dog genome for visualization in chromosomes. Peaks of high differentiation are detected despite low genome-wide divergence. Nearby annotated genes are shown above each peak. Genes with GO terms suggestive of functions relevant to salinity (*SLCA1, AKAP6*) or temperature (*DHDDS, BCL11B*) adaptation are marked in bold. A cutoff of ΔAF=0.55 is indicated. B-C) Allele frequency differences and gene annotations within 1Mbp around the highest peak regions on contigs *H. fasciata* 20 (Hfa020) (B) and 27 (Hfa027) (C).

Next, we analyzed the gene functions within genomic regions surrounding localized peaks of high ΔAF. Given the strong environmental differences between the Baltic Sea and the NE Atlantic, we hypothesized that these regions might include genes implicated in responses to habitat-specific ecological pressures, particularly salinity and temperature (osmo- and thermoregulation). Several genes had GO terms related to ion homeostasis and transport, including *SLC30A2, ATP1A3, KCNQ3* and *SLC9A1* (Table S2, S3). Genes with GO annotations related to lipid binding and metabolism (e.g., *RBP1, HSD3B7, PAFAH2*) and genes with previously described roles in thermoregulation, such as *GPR3* (Sveidahl Johansen et al. 2021) and *GADD45A* (You et al. 2020), were also detected in the vicinity of high-ΔAF regions (Table S2, S3).

We annotated the variants segregating between the subspecies using SnpEff-predicted functional effects and Genomic Evolutionary Rate Profiling (GERP) scores. The latter quantify evolutionary conservation as the reduction in the number of substitutions relative to the neutral expectations in a multi-species alignment (Huber et al. 2020). We detected no non-synonymous variants segregating between the Baltic and the Atlantic within the set of candidate genes (Table S3). However, missense variants with high ΔAF were found in a small number of other genes, such as *DSG2* and *HUWE1* (Table S2). Overall, most of the variants in the regions of high ΔAF were classified as MODIFIERS, a category that includes variants in intergenic regions, as well as introns and UTRs. Annotation with GERP scores indicated that some of these alleles, although not resulting in protein changes, are conserved across mammals (Table 2), consistent with a potential functional role in gene regulation (Li et al., 1999). In most of those cases, the derived allele had higher frequency in the Baltic (Table 2). Alignment of the dog and ribbon seal sequences further revealed that sites with GERP≥2 within the highest ΔAF peak on contig Hfa020 are highly conserved (Figure S4).

**Table 2.**
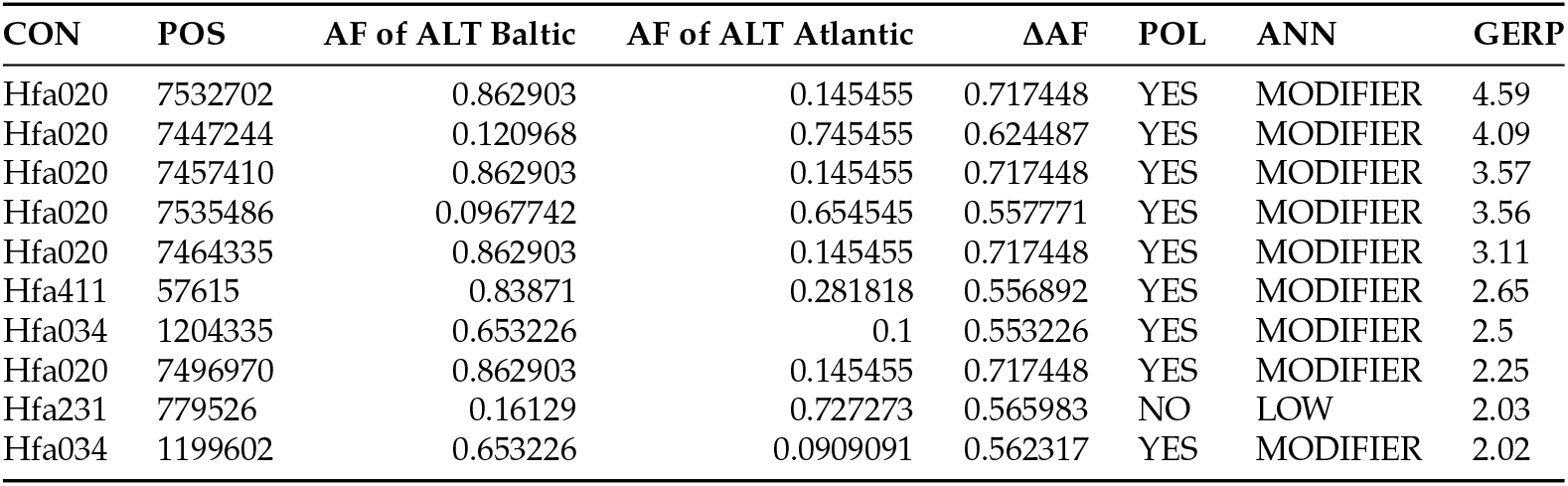
Variants with ΔAF≥0.55 between the Baltic and the Atlantic and GERP conservation scores ≥ 2. The table indicates whether the variant could be polarized (in which case the ALT allele confidently corresponds to a derived mutation) and provides the variant effect annotation. Here, both biallelic indels and biallelic SNPs are included.

| CON | POS | AF of ALT Baltic | AF of ALT Atlantic | $\Delta AF$ | POL | ANN | GERP |
| --- | --- | --- | --- | --- | --- | --- | --- |
| Hfa020 | 7532702 | 0.862903 | 0.145455 | 0.717448 | YES | MODIFIER | 4.59 |
| Hfa020 | 7447244 | 0.120968 | 0.745455 | 0.624487 | YES | MODIFIER | 4.09 |
| Hfa020 | 7457410 | 0.862903 | 0.145455 | 0.717448 | YES | MODIFIER | 3.57 |
| Hfa020 | 7535486 | 0.0967742 | 0.654545 | 0.557771 | YES | MODIFIER | 3.56 |
| Hfa020 | 7464335 | 0.862903 | 0.145455 | 0.717448 | YES | MODIFIER | 3.11 |
| Hfa411 | 57615 | 0.83871 | 0.281818 | 0.556892 | YES | MODIFIER | 2.65 |
| Hfa034 | 1204335 | 0.653226 | 0.1 | 0.553226 | YES | MODIFIER | 2.5 |
| Hfa020 | 7496970 | 0.862903 | 0.145455 | 0.717448 | YES | MODIFIER | 2.25 |
| Hfa231 | 779526 | 0.16129 | 0.727273 | 0.565983 | NO | LOW | 2.03 |
| Hfa034 | 1199602 | 0.653226 | 0.0909091 | 0.562317 | YES | MODIFIER | 2.02 |

Together, these results suggest that localized peaks of high ΔAF are in the vicinity of genes with potential functional relevance for environmental adaptations of Baltic and NE Atlantic grey seals, including genes implicated in ion transport and homeostasis (salinity), as well as lipid metabolism and thermoregulation (temperature). Although no coding variants were found within the main candidate genes, many highly differentiated non-coding variants occurred at conserved sites and could therefore have a gene regulatory function.

## 3. Discussion

In this study, we use whole-genome data to resolve the demographic history, secondary contact, and genomic differentiation of NE Atlantic and Baltic grey seals. Our results support that divergence between the subspecies was gradual, with declining gene flow broadly coinciding with the postglacial development of the Baltic Sea and the emergence of distinct Baltic environmental conditions. We confirm recent secondary contact by identifying a likely first-generation hybrid at Rødsand and a Baltic-like migrant in the North Sea, and we use sex-linked variation to infer a Baltic maternal and NE Atlantic paternal origin for the hybrid. Finally, we identify localized peaks of high differentiation that are potentially associated with non-coding regions in the vicinity of genes involved in ion transport, homeostasis, lipid metabolism, and thermoregulation, which might contribute to local adaptation of the respective sub-species.

### 3.1 Population divergence

Our WGS data allowed for more detailed insights into the process of divergence between the Baltic and Atlantic grey seals than previously possible (Graves et al. 2009; Klimova et al. 2014; Fietz et al. 2016; McCarthy et al. 2025). We found a reduction in gene flow starting 10,000 years ago, cumulative gene flow dropping below 50% 6000 years ago, and gene flow ceasing completely about 2000 years ago. The initial decline in gene flow broadly coincides with the final retreat of the Weichselian ice sheet and the early postglacial development of the Baltic Basin, when changing marine connectivity, salinity, and habitat availability likely reshaped grey seal distributions. The 50% cumulative migration estimate around 6000 years ago falls within the mid-Holocene, after the establishment of the brackish Littorina Sea phase and the development of more modern Baltic-like salinity conditions (Björck 1995; Andrén et al. 2011). Together, these timings are consistent with the geological and zooarchaeological records, which suggest that the formation of the Baltic Sea and its distinct ecological conditions may have contributed to divergence between Baltic and NE Atlantic grey seals (Ahlgren et al. 2022). By contrast, the inferred complete cessation of gene-flow about 2000 years ago is surprising as it predates the 16th-19th century local extinctions of grey seals at geographical “bridge” localities in the southern North Sea, Wadden Sea and inner Danish Waters (Prummel and Heinrich 2005; Olsen et al. 2018). However, this timing broadly coincides with the Iron Age in northern Europe and follows millennia of early coastal human exploitation, including seal hunting from the Stone Age onward (Ahlgren et al. 2022). Human impacts may therefore have already contributed to reduced connectivity between Baltic and NE Atlantic grey seals, although this interpretation remains speculative. Several caveats should be also considered when interpreting these divergence-time estimates. First, the estimates depend on the demographic parameters used in the model, which may not be optimal for grey seals. Second, they are inferred from present-day populations and may not capture the genetic composition of historical populations that once inhabited the North Sea region. Indeed, current levels of connectivity in the North Sea region might be a recent phenomenon driven by source-sink metapopulation dynamics, whereas genetic and demographic connectivity may have been lower in the past (McCarthy et al. 2025).

### 3.2 Overall population structure and hybridization

As in McCarthy et al. (2025), we found the NE Atlantic and Baltic subspecies to be genetically distinct. The NE Atlantic population has a more pronounced stratification, with distinct populations in the North Sea region, Iceland and Norway-Russia. Interestingly, our analyses also pointed to some substructure within the Baltic Sea, with grey seals at the Danish Rødsand haul-out site clustering distinctly from those sampled in Sweden, Estonia and Finland and showing high genetic similarity within the site. This genetic signature could be a bias caused by Rødsand animals on average being more related (Table S1), perhaps indicating site-fidelity and the early stages of (re-)establishment of a breeding colony at Rødsand highly influenced by a limited number of ‘founder’ individuals. Alternatively, one might tentatively hypothesize that the genetic distinctiveness of Rødsand grey seals reflects a relict genomic component that somehow survived the (presumed) 19th century local extinctions in the region.

We confirm the existence of ongoing hybridization by providing clear evidence for a likely first-generation hybrid. The hybrid animal was a pup sampled at the Rødsand haul-out site in the Danish part of the SW Baltic in 2013 (Table S4), genetically inferred to have a Baltic Sea mother and a NE Atlantic father. Previous studies have hinted at the presence of admixed individuals at that site (Fietz et al. 2016; McCarthy et al. 2025); however this is the first confirmed detection of a hybrid. In total, we sampled and sequenced 20 pups from Rødsand, representing approximately 22% of all the pups born there during the sampling period, and all other pups were genetically identified to have Baltic parental heritage. This indicates that hybridization might still be rare. In their core range, NE Atlantic and Baltic grey seal are characterized by marked differences in the timing of their pupping seasons (Härkönen et al. 2007; Galatius et al. 2024), which may take time to synchronize in the intermediate region of recovery and recolonization. Nevertheless, the species is known for its plasticity in timing of breeding (Bowen et al. 2020), and it is therefore possible that males are able to mate outside their normal breeding season.

### 3.3 Localized genomic differentiation and potential local adaptation

The regions surrounding the identified peaks that show high ΔAF between the subspecies included multiple gene annotations with GO terms suggesting functions in ion and osmoregulation or lipid metabolism and thermoregulation. For example, *SLC30A2, ATP1A3, KCNQ3* and Na^+^/H^+^ exchanger genes such as *SLC9A1* have been implicated in salinity and osmotic regulation in fish (Liu et al. 2019; Wen et al. 2021; Han et al. 2025). *GPR3* has been found to activate mouse and human thermogenic adipocytes (Sveidahl Johansen et al. 2021), while *GADD45A* has been associated with non-shivering thermogenesis in mice (You et al. 2020). We hypothesized that such functions may be relevant to differential salinity and/or cold adaptation in the two subspecies, with Baltic Sea seals inhabiting a region characterized by strong salinity gradients and marked seasonal variation in air and sea temperatures, including the presence of winter sea-ice (Stramska and Białogrodzka 2015). Notably, adaptive genetic changes related to osmoregulation have been reported in *Pusa* seals inhabiting low-salinity environments (Zhao et al. 2026), along with potentially adaptive changes in kidney anatomy (Nihtilä and Laakkonen 2025).

Although these functional annotations are suggestive, only a subset of the genes near ΔAF peaks is likely to be directly involved in local adaptation. We found no coding variants within the main candidate genes, indicating that adaptive divergence, if present, may more likely involve regulatory rather than protein-coding changes. Putative regulatory non-coding variants are harder to identify, as *cis*-regulatory elements may act from great distances upstream or downstream of genes (Kim and Wysocka 2023), or from intronic regions (e.g., Kratochwil et al., 2018). Nevertheless, segregating variants with high GERP scores in the highest peak region on contig Hfa020 and in other regions (Table 2) are likely to have a regulatory role, altering gene expression without requiring fixed amino-acid differences. This is plausible given the recent divergence and recolonization history of grey seals, their high dispersal potential, and the strong seasonal and interannual fluctuations in salinity and temperature across their habitats. Moreover, genes with a less obvious connection to salinity or cold adaptation might still be relevant. For example, the missense mutation inside *DSG2* (desmoglein-2 gene) may be of interest, as genes coding for components of the desmosome have been found to be positively selected in marine mammals (Chikina et al. 2016), potentially enhancing the resistance of the skin and connective tissues to heat loss or osmotic stress. The observation that derived mutations in conserved sites were more frequent in the Baltic subspecies (Table 2) is consistent with the Atlantic subspecies being the ancestral population and colonizing populations into the Baltic Sea having adapted to the unique environment. In any case, such localized genetic differences between the two subspecies are notable given their recent history of divergence.

### 3.4 Possible long term evolutionary consequences

As the overlap between the NE Atlantic and the Baltic grey seal populations is still in early stages (Fietz et al. 2016; Galatius et al. 2024), predictions about the evolutionary consequences of hybridization in this system remain speculative. Despite their recent divergence, the populations have differentiated across several genomic regions, potentially reflecting differences in selective pressures. While the hybridization process unfolds, it is possible that a subset of these regions will have fitness consequences in hybrid individuals and thereby limit introgression by acting as barriers to the genetic homogenization of the NE Atlantic and Baltic grey seals (Feder et al. 2013; Harrison and Larson 2016).

Environmental gradients and transitions, such as the changes in salinity and temperature conditions between the Atlantic and the Baltic Sea have been shown to influence hybridization dynamics in many systems (e.g., Culumber et al., 2012; Haenel et al., 2021). Whether, in this case, the differences between the subspecies will eventually form a continuous gradient, as previously hypothesized (McCarthy et al. 2025), or a steeper shift as observed in other Atlantic-Baltic population pairs (Johannesson and André 2006), remains to be explored. Throughout this process, the exchange of adaptive alleles is also a possibility. Given that “the pattern of contemporary hybridization is potentially only a single snapshot of a complex and continuously changing interaction” (Abbott et al. 2013), continuous monitoring and research will be necessary to investigate the outcomes of the hybridization between the grey seal subspecies and its implications for the species’ recolonization and recovery.

### 3.5 Conclusions

The recolonization process of grey seals following recovery from human-induced range restrictions and local extinctions has resulted in secondary contact, and thus the potential for hybridization, between the NE Atlantic and the Baltic subspecies (Fietz et al. 2016; Galatius et al. 2024). Using whole-genome sequencing data, we estimated that the Baltic and the Atlantic subspecies diverged approximately 6000 years ago, identified a Baltic migrant into the NE Atlantic, and confirmed hybridization between subspecies in the form of a single hybrid individual, sampled in the SW Baltic, with inferred Baltic maternal and Atlantic paternal ancestry. The genomes of the subspecies show localized regions of high differentiation, potentially driven by differential adaptation to Atlantic and Baltic environmental conditions. Our findings emphasize the evolutionary relevance of contemporary hybridization, placing it in the context of population recolonization and recovery, and call for further research into how evolutionary dynamics will unfold in the case of grey seals and in other systems recovering from human disturbances or facing new pressures due to the ongoing climate and biodiversity crises.

## 4. Materials and Methods

### 4.1 Sampling, DNA extraction and whole-genome sequencing

Grey seal tissue samples were collected across the northeastern distributional range of the species (Figure 1A; Table 1; Table S4 in detail), including 20 grey seal pups that were sampled in the period 2013-2025 at the Rødsand haul-out site in SW Baltic to specifically match the time and space of increasing distributional overlap between the two subspecies. These 20 pups represent approximately 22% of the pups recorded at the site under the Danish monitoring program during this period. Sample collection was done from live animals under permits from national authorities, from animals by-caught in fishing gear, from hunted or from dead beach-cast animals recovered through national stranding networks. Samples were stored at −20°C until DNA extraction, which was done following the manufacturer’s instructions using the Qiagen DNeasy Blood and Tissue Kit, the Thermo Fisher Scientific KingFisher Cell and Tissue DNA Kit or the MagMAX™ DNA Multi-Sample Ultra 2.0 Kit. The quantity and quality of the extracted genomic DNA were assessed on a Qubit Fluorometer and using 1% agarose gels with 1 kb ladders, respectively. Whole-genome sequencing of genomic DNA was completed by BGI Genomics using PCR-free library preparation and PE 150bp DNBseq with a few individuals per sampling site sequenced at high coverage (>30x) and all remaining samples at medium coverage (10-20x).

### 4.2 Positive mask and functional categories

Genomic regions where short sequence reads cannot be unambiguously mapped were identified following Olkkonen and Löytynoja (2023). The repeat regions, covering 42.2% (1.027 Gbp) of the used ribbon seal (*Histriophoca fasciata*) reference genome, were identified with RepeatMasker (v.open.4.05) and RMBlast (v.2.6.0) (Smit et al. 1996) using dog repeat libraries. The mappable regions, covering 90.9% (2.213 Gbp) of the reference genome, were identified with SNPable (seqbility-20091110; Li, 2009) using 75 bp fragments. We combined the mapping and repeat masks using BEDtools (v.2.30.0; Quinlan and Hall, 2010), resulting in a positive mask of 1.328 Gbp, corresponding to 54.5% of the reference genome.

Functional regions were annotated using information from related species. The protein sequences of the elephant seal genome annotation (v.GCF_029215605.1-RS_2024_04) (NCBI 2024) were mapped to the reference genome with miniprot (v.0.13-r262-dirty; Li, 2023), specifying the option–gff to produce the GFF-formatted annotation file. To remove spurious hits, only transcripts with sequence identity of 80% or greater were retained, reducing the number of transcripts from 64,643 to 63,946. BUSCO (v.5.8.2, odb12; Manni et al., 2021) analysis was performed with the lineage “carnivora”, and the miniprot-created GFF file of BUSCO genes was extracted. The intersection of elephant seal CDSs and BUSCO CDSs was computed with BEDtools, requiring reciprocal 90% coverage. To obtain corresponding Ensembl gene IDs and gene symbols for downstream analyses, we aligned the elephant seal proteins against ferret protein sequences (MusPutFur1.0.pep.all; Ensembl version 115; Dyer et al., 2025) using BLAST (v.2.16.0; Camacho et al., 2009) and, for each, retained the first match with identity of at least 75% and *E*-value < 1×10^-7^ .

### 4.3 WGS data processing

The WGS dataset used in this study consisted of 123 individuals, of which 119 were retained for analysis (56 NE Atlantic; 63 Baltic; Table 1). The four excluded samples showed high coverage variance and extensive regions of low coverage (Figure S5A), which appeared to bias population structure and genetic distance inference (Figure S5B,C). This dataset was supplemented with read data from 17 Arctic ringed seal (*Pusa hispida*) individuals from Greenland (Table 1), described in Löytynoja et al. (2023), to serve as an outgroup in the *f* -statistics estimation.

The FASTQ files were mapped to a ribbon seal reference genome with BWA MEM2 (v.2.2.1; Vasimuddin et al., 2019), and regions around indels were realigned with IndelRealigner from Genome Analysis Toolkit (GATK: GATK3 v.3.8-1-0-gf15c1c3ef; McKenna et al., 2010). Variants were called with HaplotypeCaller from GATK4 (v.4.5.0; McKenna et al., 2010) and SAMtools (v.1.16.1; Danecek et al., 2021). GVCFs were combined with GenomicsDBImport and jointly genotyped using GenotypeGVCFs. To generate an outgroup for computing *f* -statistics, we additionally performed joint calling of existing GVCF files from Greenlandic Arctic ringed seals, restricting it to sites included in the grey seal VCF (options “-L $VCF –force-output-intervals $VCF”). The VCFs were merged with BCFtools merge (v.1.16.1; Danecek et al., 2021).

The VCF file was filtered to include only biallelic SNPs that are within positively masked regions, segregate among the samples and show sequencing depth between 1000 and 3500 (total across individuals). We also created a version of the VCF with stricter depth filtering (1400<DP<3500) to use in the Identity-by-state distance estimation, which has been shown to be particularly sensitive to low-coverage data (Olkkonen and Löytynoja 2023). All filtering steps were done with BCFtools view.

As a quality check, we estimated the sequence depth of the largest contig (Hfa001; 64.772 Mbp) in 50 kbp windows in individual BAM files using SAMtools depth. Coverage distributions of all grey seal samples were inspected in R (v.4.5.2; R Core Team 2024). The mean coverage per individual, averaged across windows, ranged between 10.16X and 57.66X, and the overall mean was 18.53X.

To infer relatedness among all individuals, and particularly to investigate the structure observed in the Rødsand population (see 2.2), we used the KING toolset (v.2.3.2; Manichaikul et al., 2010) with the “–related” option (Table S1).

### 4.4 Demographic and isolation-migration inference

MSMC2 (Schiffels and Wang 2020) and MSMC-IM (Wang et al. 2020) were applied to infer demographic history, migration rates, and population split times. First, the full data were imputed and phased with Beagle (v.4.1; Browning et al., 2021) for Baltic (n=34) and Atlantic grey seals (n=28; excluding Norway and Russia, see Figure 2A). *N*_e_ was specified for imputation and phasing (Pook et al. 2020) as: *N*_e_ = 50,000 for Baltic and *N*_e_ = 100,000 for Atlantic grey seals. Imputed and phased VCF files were merged with BCFtools merge. Six samples with the highest sequencing depth were selected from each group, covering all sampling locations. 414 contigs that were at least 250 kbp long and contained at least 1000 SNPs were included in the analysis. MultiHetSep files for positively-masked regions in these contigs were generated using generate_multihetsep.py from the MSMC-tools package (Schiffels and Wang 2020).

For the demographic inference, MSMC2 analyses were performed with four individuals per analysis (e.g., -I 0,1,2,3,4,5,6,7) with a customized segmentation; i.e., the first two time windows were split (-p 27*1+1*2+1*3), as recommended by Hilgers et al. (2025) to avoid false recent *N*_e_ peaks. This resulted in 15 MSMC2 analyses per group. The time boundaries and coalescence rates were converted into years and effective population size, using mutation rate of μ = 1.829×10^-8^ from the polar bear (Liu et al. 2014) and a generation time of g = 10 years as in Löytynoja et al. (2023).

To derive population split times and migration patterns, a continuous Isolation-Migration (IM) model (Wang et al. 2020) was fitted onto the MSMC2 rCCR outputs. First, MSMC2 analyses with two samples per analysis (e.g., -I 0,1,2,3) and customized segmentation were performed. Each of the six samples was combined with every other sample in the group, resulting in 15 MSMC2 analyses per group. Second, MSMC2 analyses between two individuals from each subspecies (e.g., -I 0-4, 0-5, 0-6, 0-7, 1-4, 1-5, 1-6, 1-7, 2-4, 2-5, 2-6, 2-7, 3-4, 3-5, 3-6, 3-7), were performed, resulting in 25 analyses. Third, the MSMC2 analysis output files were combined using combineCrossCoal.py from the MSMC-tools package. Finally, MSMC_IM.py from the MSMC-IM-tools package was applied to the combined output files with the settings β = 1e-8,1e-6, as recommended by Wang et al. (2020), and μ = 1.829×10^-8^ (Liu et al., 2014). The time boundaries were converted into years, and the migration rate was cut at M=0.999 in R using the package dplyr (Wickham et al. 2025). Results were visualized using ggplot2 (Wickham 2016).

### 4.5 Genetic differentiation and population structure

PCA, IBS distance estimation and admixture analysis were used to infer population structure, genetic distances and possible signals of admixture. For the PCA and the admixture analysis we used a thinned version of the grey seal VCF, containing 1,594,645 SNPs, to improve computational efficiency. Thinning was done with PLINK (v.1.90b7.7; Purcell et al. 2007), applying a 1-kbp minimum distance filter between SNPs. The PCA was performed with smartpca (Price et al. 2006) from the EIGENSOFT (v.7.2.1; Patterson et al. 2006) software package. The normalization option was set to ‘NO’. For visualization we used the R packages ggplot2, ggrepel (Slowikowski 2024) and wesanderson (Ram and Wickham 2023). IBS distances were estimated with PLINK using the VCF file with stricter depth filtering and the options “–distance 1-ibs square gz –allow-extra-chr”. We used dplyr and tidyr (Wickham et al. 2026) for data manipulation in R and pheatmap (Kolde 2025) and wesanderson for visualization. For the admixture analysis we used ADMIXTURE (v.1.3.0; Alexander et al., 2009) and visualized the results for *K*=2-5 using ggplot2, wesanderson and cowplot (Wilke 2025).

### 4.6 Assessment of hybrid status of HG301

The genetic clustering approaches identified one individual, HG301, as a possible hybrid between the Baltic and the Atlantic subspecies. To verify this and quantify the proportion of each ancestry, we estimated the *f*_*4*_-statistic and implemented the *f*_4_-ratio test from the ADMIXTOOLS (v.2.0.4; Maier and Patterson, 2024) R package. The *f*_4_-statistic is computed in the form (A,B;C,D), which denotes four populations, and measures the covariance of allele frequency difference between the pairs A-B and C-D across loci. When D represents an outgroup population, then a significantly negative statistic indicates gene flow between C and B. To test for gene flow both from the Baltic and the NE Atlantic into HG301 we used the topologies [UK, HG301; (SWE_STA+FIN+EST), GRE_RIN] and [(SWE_STA+FIN+EST), HG301; UK, GRE_RIN], respectively (see Figure 1A for locality information). Here UK corresponds to the pooled UK populations, SWE_STA+FIN+EST to the merged Stockholm Archipelago, Finnish and Estonian populations and GRE_RIN includes all 17 Greenlandic ringed seal individuals that were used as an outgroup. The *f*_4_-statistic was additionally computed to confirm the absence of admixture from both subspecies into other individuals and populations. *f*_4_-ratio tests were then used to estimate the proportion of ancestry (α) that HG301 derived from the Baltic and Atlantic subspecies. SWE_FAL and UK were set as lineage B to compute α from the Baltic and the Atlantic, respectively (Figure S3). Lineage A is assumed to be a Baltic (first case) or an Atlantic (second case) population independent of B. It should be noted that this independence assumption is not fully met for the Baltic, as previous analyses did not show structure within the subspecies.

### 4.7 Sex-linked contigs and sex identification

Contigs belonging to the X chromosome were identified using three complementary approaches: coverage ratio comparison between males and females (m:f), heterozygosity comparison and homology to the dog (*Canis familiaris*; ROS_Cfam_1.0.dna_sm.toplevel; Ensembl version 115) X chromosome and the Saimaa ringed seal (*Pusa saimensis*; Sundell et al., 2023) X-linked scaffolds. As males are the heterogametic sex, they are expected to have half of the female coverage and heterozygosity values near zero across contigs belonging to the X chromosome.

Coverage in 10-kbp windows was estimated with deepTools (v.3.5.4; Ramírez et al., 2014), applying RPKM (Reads Per Kilobase Million) normalization, in 10 samples with available sex information (5 per sex). The coverage of each window was normalized to the average coverage of the five largest autosomal contigs (here, Hfa001-Hfa005), which showed no notable coverage differences between individuals. Next, we averaged the coverage of windows within each contig and tested for significant differences between sexes using a non-parametric Wilcoxon Rank Sum Test with a Bonferroni multiple test correction, following Catalán et al. (2025). From the contigs that showed significant differences, the ones with a m:f coverage ratio of 0.5±0.1 were selected. One candidate X contig, Hfa209, contained the *TSPYL2* (testis-specific Y-encoded-like protein 2) gene, which in dogs and humans is located on the X chromosome (Ensembl release 115). Males and females in the full dataset were separated based on normalized coverage values (averaged across windows) on this contig. As a sanity check, we estimated the sex ratio of the dataset, which in total contained more males (55%) than females (45%).

Heterozygosity (*H*) in windows for all individuals was computed following Olkkonen and Löytynoja (2023). First, genetic diversity of individuals was estimated per site with VCFtools –site-pi (v.0.1.17; Danecek et al., 2011). Using these estimates, *H* in 250 kbp genomic windows was calculated in R using the packages data.table (Barrett et al. 2026), IRanges (Lawrence et al. 2013), scales (Wickham, Pedersen, et al. 2025), and tidyverse (Wickham et al. 2019), correcting the window width with the positive mask and nan-sites. For the correction, missing data were assumed to be randomly distributed, and the total lengths were corrected as (*S*−*n*)/*S*×*L*, where *S* and *n* are the numbers of variant sites and nan sites, and *L* is the length of the subset after applying the positive mask. Then, using the inferred sex assignments, a Wilcoxon Rank Sum Test was performed to detect contigs with significant differences in *H* between males and females. Finally, we mapped the ribbon seal reference genome against a dog and a Saimaa ringed seal genome, using minimap2 (v.2.28; Li, 2018) to identify contigs aligning to the dog X chromosome and the *P. saimensis* X scaffolds, and compiled a final set of X contigs.

### 4.8 The hybrid’s parental populations

Since the hybrid individual was inferred to be a male, containing only one X chromosome copy, its maternal population could be determined by performing a PCA using only X-linked contigs and only male samples, haploid for those contigs. It is expected that, if the hybrid’s X contigs cluster with males of one subspecies, then the X chromosome and hence its mother derives from that subspecies, implying that its father belongs to the other subspecies. We performed the PCA with smartpca from EIGENSOFT and visualized the results with ggplot2, ggrepel and wesanderson.

### 4.9 Allele frequency estimation and gene functions

To explore genomic differentiation between the Baltic and the Atlantic populations, allele frequency difference (ΔAF) was estimated across sites. Allele frequencies per site were first computed for the Baltic and the Atlantic individuals separately, using VCFtools (option “–freq”), and then absolute frequency difference of one of the alleles in each position (e.g., reference allele) between populations was calculated in R. To visualize ΔAF in chromosomes, we lifted the values over to a dog genome (ROS_Cfam_1.0.dna_sm.toplevel). For this, a liftover chain was generated between the ribbon seal reference and the dog genome using the Nextflow LiftOver pipeline (Talenti and Prendergast 2021) and then the ΔAF values were transferred to the corresponding coordinates of the dog genome using swiftover (Blachly lab 2023). The R package qqman (Turner 2018) was used to plot genome-wide ΔAF.

To inspect genes in proximity to the areas of high differentiation, the annotation of functional regions described above was used. For visualizing genes surrounding the highest ΔAF regions we used the R package Gviz (v.1.51.0; Hahne and Ivanek, 2016). To detect genes potentially related in cold or salinity adaptation, we extracted all genes within 1 Mbp of regions of high allele frequency difference (Table S2) and filtered them based on Gene Ontology (GO) terms, retaining those with annotations related to ion and osmotic regulation or lipid metabolism (e.g., containing “osmo”, “sodium”, “potassium”, “chloride”, “anion”, “cation”, “lipo”, “lipid”). For this, GO annotations for ferret gene IDs (MusPutFur1.0) were retrieved from Ensembl using the R package biomaRt (Durinck et al. 2009). We made minor refinements to the resulting gene list based on a literature search (Table S3).

### 4.10 Variant effect prediction and conservation scores

SnpEff (Cingolani et al. 2012) was used to predict the functional effect of variants segregating between the subspecies. Prior to this, we inferred the ancestral state of alleles and repolarized a version of the VCF without positive mask filtering and including both biallelic SNPs and biallelic indels. For that, HiFi data from the elephant seal (SRR25478317_subreads.fastq.gz) were aligned to the reference genome with minimap2 (options “-a-x map-hifi”). When inferring the ancestral state for positions in a VCF file, the homologous position in the elephant seal was called with GATK HaplotypeCaller (options “-L $VCF – output-mode EMIT_ALL_CONFIDENT_SITES-ERC BP_RESOLUTION”) and GenotypeGVCFs (options “-L $VCF–call-genotypes true–force-output-intervals $VCF”). The ancestral states were resolved with a custom script that requires read depth >4 and the homozygous allele in the elephant seal to exactly match either the reference allele or the alternative allele in the input VCF data. The AA tag was added in the VCF file with BCFtools annotate, and at positions where AA matches ALT, the VCF data were repolarized following Abascal (2016).

To assess whether non-coding variants with high ΔAF might be functionally important, possibly belonging to regulatory regions, this VCF was also annotated with GERP scores obtained from Ensembl datasets for ferret (https://ftp.ensembl.org/pub/release-115/compara/conservation_scores/92_mammals.gerp_conservation_score/). For lifting the GERP scores from the ferret to the ribbon seal genome, we used the Nextflow LiftOver pipeline to generate the ferret-to-seal chain file and swiftover to convert the coordinates. Then, the VCF was annotated with the GERP score tag using BCFtools annotate. To further inspect conservation patterns around the highest ΔAF peak, we aligned it to its homologous region of the dog genome using the Shuffle-LAGAN alignment method (Brudno et al. 2003) from the mVISTA server. For this, the sequence 800kbp upstream and downstream of the peak was extracted from the ribbon seal genome using SAMtools faidx, as well as the corresponding region of the dog genome. Gene annotations for the dog genome, extracted from Ensembl in VISTA format, were also included.

## Acknowledgements

MLM, IS and MTO were supported by a Carlsberg Foundation Semper Ardens Accelerate fellowship to MTO (grant CF21-0425). CFK was supported by a Research Council of Finland Fellowship and Research Grant (347309, 371328) and a Sigrid-Jusélius Foundation Grant.

## Author contributions

AK, AL, AG, CFK and MTO conceived and designed the research; AG, SG, BM, MJ, IJ, MK, US, AH, PA, JJ, RD, JT and MTO provided samples and funding; IS and MTO performed the laboratory analyses; AK, AL, TK and MLM analyzed the data; AK drafted the manuscript together with AL, CFK and MTO; all authors provided editorial inputs and approved the final version of the manuscript.

## Data availability

Short read sequences will be deposited to the European Nucleotide Archive (ENA) upon acceptance of the manuscript.

Documentation of the code used in this study is available at https://github.com/AnastasiaKonstantopoulou/seal_hybridization_paper.

## Notes

### Competing Interest Statement

The authors have declared no competing interest.

