## Supplementary Figures 1-5; Supplementary Tables 1-4 for "Contemporary hybridization and localized genomic differentiation between grey seal subspecies"

[1]Faculty of Biological and Environmental Sciences, University of Helsinki, Finland
[2]Institute of Biotechnology, Helsinki Institute of Life Science (HiLIFE), University of Helsinki, Finland
[3]Department of Computer Science, University of Helsinki, Finland
[4]Marine Mammal Research, Department of Ecoscience, Aarhus University, Roskilde, Denmark
[5]Section of Molecular Ecology and Evolution, Globe Institute, University of Copenhagen, Denmark
[6]Department of Population Analysis and Monitoring, Swedish Museum of Natural History, Stockholm, Sweden
[7]Section for Veterinary Clinical Microbiology, Department for Veterinary and Animal Sciences, University of Copenhagen, Denmark
[8]The Icelandic Seal Center, Hvammstangi, Iceland
[9]Marine and Freshwater Research Institute, Hafnarfjörður, Iceland
[10]Department of Biology, Norwegian University of Science and Technology, Trondheim, Norway
[11]Pro Mare, Estonia
[12]Department of Biology, University of Eastern Finland, Joensuu, Finland
[13]Natural Resources Institute Finland, Joensuu, Finland
[14]Leibniz Institut for Zoo and Wildlife Research, Berlin, Germany
[15]Sea Mammal Research Unit, Scottish Oceans Institute, University of St. Andrews, UK
[16]Department of Geosciences and Geography, University of Helsinki, Finland
[*]Equal contribution; corresponding authors

### Supplementary Figures and Tables

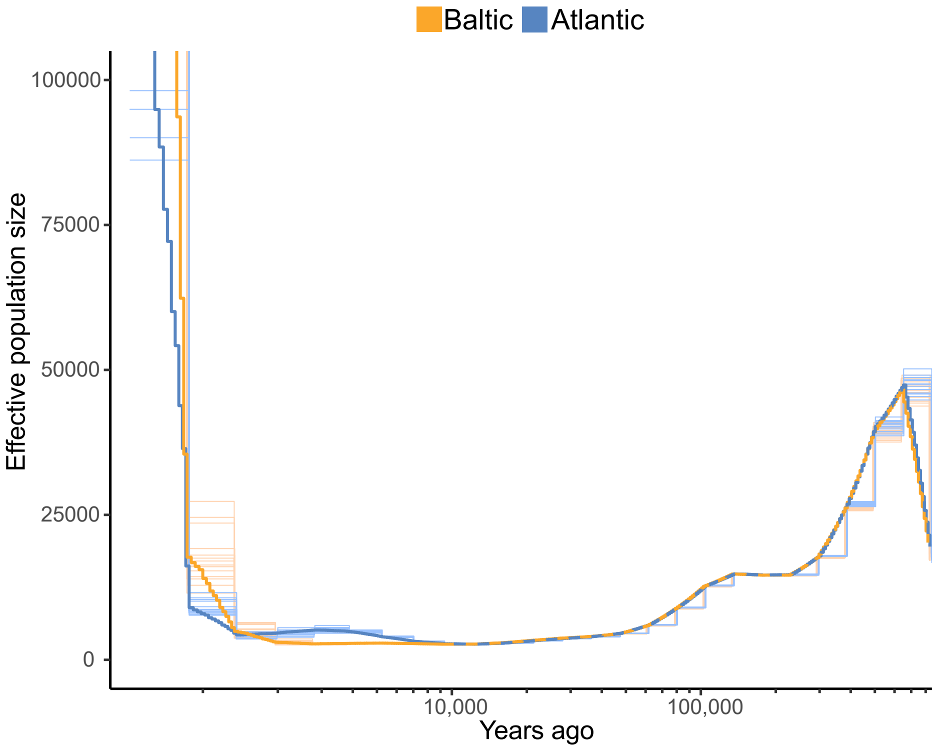


Figure S1. Estimated effective population size (Ne, y axis) for the past 800,000 years (x-axis). Individual pairs are shown with thin lines and the population averages with thick lines.

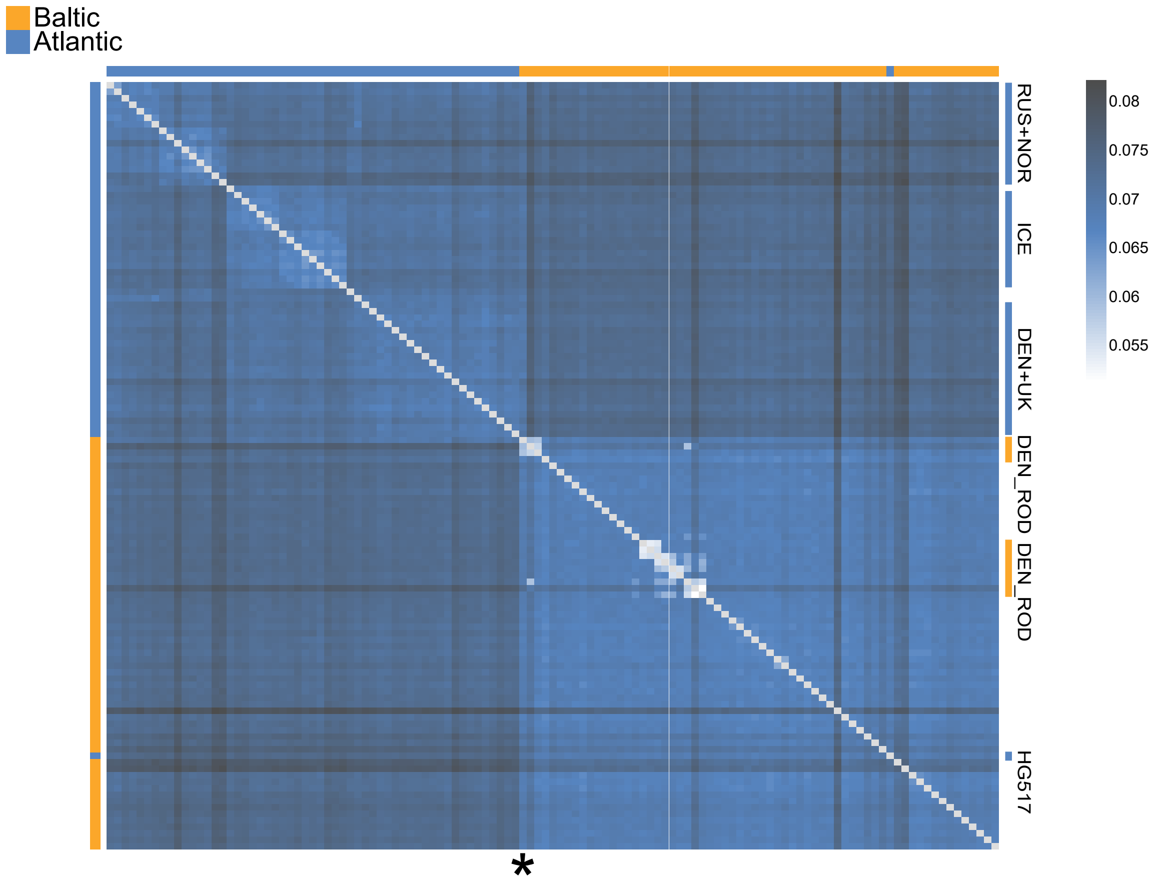


Figure S2. Heatmap of pairwise IBS distances. Darker colors indicate greater distance. Individuals are ordered based on their clustering on a neighbor-joining (NJ) tree. The identified subspecies hybrid individual, HG301, is marked with an asterisk. HG301 shows comparable genetic similarity to both Baltic and Atlantic individuals, and HG517 (noted) clusters within the Baltic.

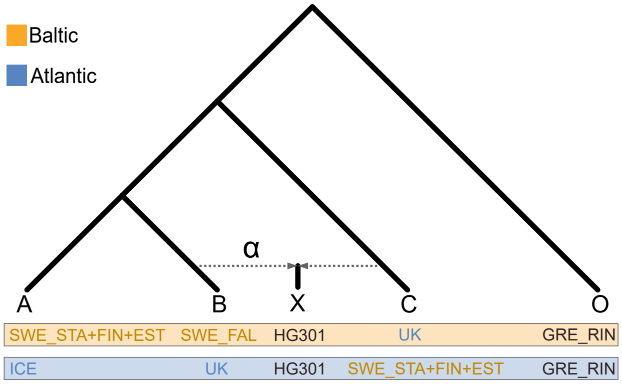


Figure S3. Setup used in the $f_4$-ratio tests to estimate the ancestry proportions ($\alpha$) coming from the Baltic (top) and the Atlantic subspecies (bottom) into HG301.

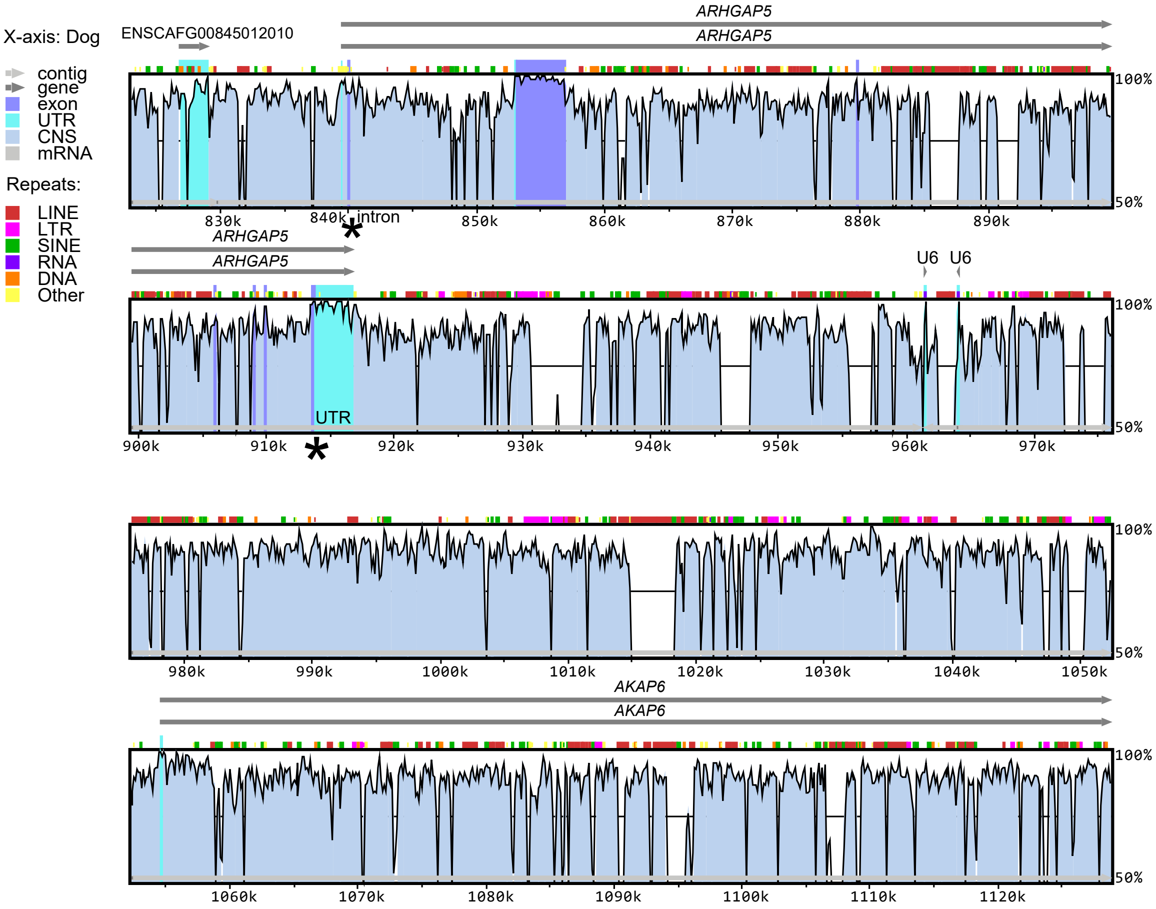


Figure S4. Conservation patterns around the highest ΔAF peak, based on the alignment of the ribbon seal reference genome to the dog genome. The x-axis shows positions in the dog genome, with corresponding gene annotations. Asterisks mark the locations of the variants with the highest GERP scores, located within an intron and a UTR of *ARHGAP5* in the dog genome.

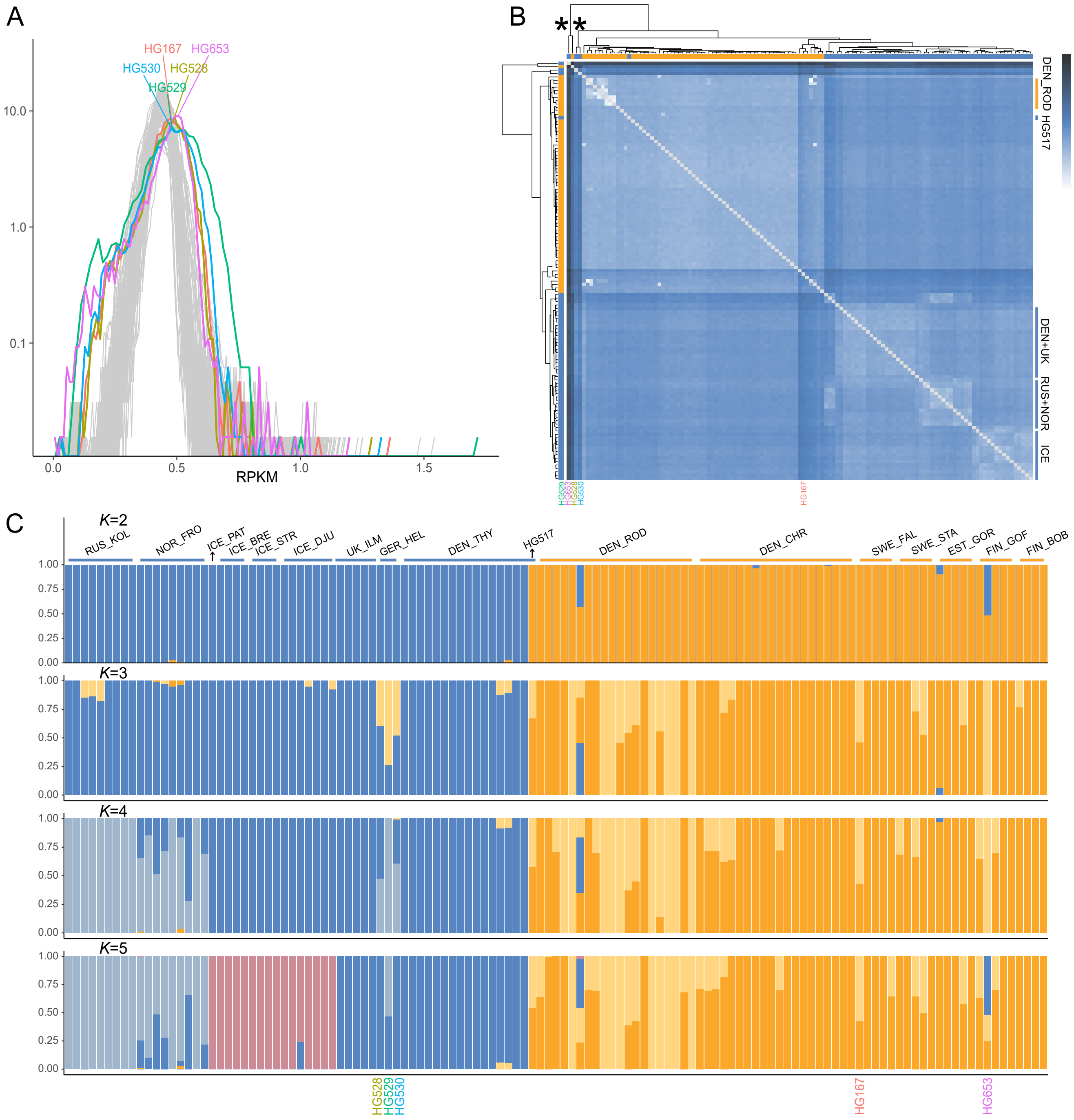


Figure S5. Data coverage distribution and its effect on population structure and genetic distance inference. A) RPKM (Reads Per Kilobase Million)-normalized coverage in 10-kbp windows across contig Hfa001 for each individual. Coverage values are summarized into evenly spaced bins. The y-axis is log-scaled. Each line represents one individual; colored lines denote samples with problematic coverage distributions (greater variance, regions of low coverage). B) Heatmap of pairwise IBS distances with clustering dendrograms. Individuals HG528, HG529, HG530 and HG653 cluster separately from all the rest (marked with two asterisks). C) Admixture analysis for $K$=2-5 genetic clusters. Individuals with problematic coverage distributions are marked. HG528, HG529, HG530 and HG653, which show the most abnormal clustering, were excluded from the main analyses.

Table S1. Relatedness inference. Pairs of related individuals identified in the dataset, all belonging to the Rødsand population (DEN_ROD). Kinship coefficient estimates and inferred degrees of relatedness are shown.

| Ind 1 | Ind 2 | IBS | Dist | Kinship | Degree |
|---|---|---|---|---|---|
| HG300 | HG304 | 1.8950 | 0.1356 | 0.0594 | 3rd-degree |
| HG300 | HG514 | 1.8824 | 0.1433 | 0.0475 | 3rd-degree |
| HG305 | HG310 | 1.8870 | 0.1363 | 0.0653 | 3rd-degree |
| HG306 | HG310 | 1.8852 | 0.1399 | 0.0502 | 3rd-degree |
| HG306 | HG311 | 1.8913 | 0.1297 | 0.0807 | 3rd-degree |
| HG307 | HG516 | 1.8877 | 0.1345 | 0.0683 | 3rd-degree |
| HG310 | HG311 | 1.8856 | 0.1382 | 0.0548 | 3rd-degree |

Table S2. Genes within 1 Mbp of regions or variants with ΔAF $\geq 0.55$

| Contig | Start | End | Dog chr | NCBI elephant seal protein ID | Ensembl ferret gene ID | Gene symbol |
|---|---|---|---|---|---|---|
| Hfa001 | 16921320 | 17424649 | 13 | XP_045737040.1 | ENSMPUG00000001363.1 | *ADGRL3* |
| Hfa001 | 45940639 | 45976313 | 3 | XP_045737330.1 | ENSMPUG00000016724.1 | *ANAPC4* |
| Hfa001 | 45985588 | 46029430 | 3 | XP_045737331.1 | ENSMPUG00000016727.1 | *ZCCHC4* |
| Hfa001 | 46045770 | 46072559 | 3 | XP_045737332.1 | ENSMPUG00000016729.1 | *PI4K2B* |
| Hfa001 | 46132786 | 46164410 | 3 | XP_045737336.1 | ENSMPUG00000016732.1 | *SEPSECS* |
| Hfa001 | 46241507 | 46269441 | 3 | XP_045737339.1 | ENSMPUG00000016733.1 | *LGI2* |
| Hfa001 | 46334851 | 46434282 | 3 | XP_045737342.1 | ENSMPUG00000016734.1 | *CCDC149* |
| Hfa001 | 46442976 | 46443719 | 3 | XP_045737343.1 | ENSMPUG00000020131.1 | *SOD3* |
| Hfa001 | 46611884 | 46668313 | 3 | XP_045737344.1 | ENSMPUG00000016735.1 | *DHX15* |
| Hfa003 | 45806576 | 45809367 | 12 | XP_064453878.1 | ENSMPUG00000018859.1 | *TPBG* |
| Hfa003 | 45912007 | 46000540 | 12 | XP_045723111.1 | ENSMPUG00000005527.1 | *IBTK* |
| Hfa003 | 46431514 | 46434180 | 12 | XP_045723107.1 | ENSMPUG00000018098.1 | *TENT5A* |
| Hfa005 | 11627255 | 11661274 | 15 | XP_064441296.1 | ENSMPUG00000001877.1 | *TMA16* |
| Hfa005 | 11674150 | 11676027 | 15 | XP_045735317.1 | ENSMPUG00000001875.1 | *TKTL2* |
| Hfa005 | 11781228 | 11782568 | 15 | XP_045736151.1 | ENSMPUG00000018775.1 | *NPY5R* |
| Hfa005 | 11787916 | 11807471 | 15 | XP_045736154.2 | ENSMPUG00000001873.1 | *NPY1R* |
| Hfa005 | 11924417 | 11962217 | 15 | XP_045736148.1 | ENSMPUG00000001861.1 | *NAF1* |
| Hfa005 | 12816151 | 13509956 | 15 | XP_045735318.1 | ENSMPUG00000016662.1 | |
| Hfa007 | 37054453 | 37115292 | 23 | XP_045759618.1 | ENSMPUG00000012312.1 | *SPSB4* |
| Hfa007 | 37394571 | 37418838 | 23 | XP_045719671.2 | ENSMPUG00000004993.1 | *TRIM42* |
| Hfa007 | 37501577 | 37637202 | 23 | XP_045759620.1 | ENSMPUG00000004999.1 | *CLSTN2* |
| Hfa007 | 38360975 | 38422733 | 23 | XP_045759622.1 | ENSMPUG00000009073.1 | *NMNAT3* |
| Hfa007 | 38439798 | 38463659 | 23 | XP_045759625.1 | ENSMPUG00000009083.1 | *RBP1* |
| Hfa007 | 38503646 | 38525870 | 23 | XP_045719672.1 | ENSMPUG00000009092.1 | *RBP2* |
| Hfa007 | 38586672 | 38611638 | 23 | XP_045759626.1 | ENSMPUG00000009098.1 | *COPB2* |
| Hfa007 | 38612961 | 38640317 | 23 | XP_045759627.2 | ENSMPUG00000009114.1 | *MRPS22* |
| Hfa007 | 39186568 | 39297343 | 23 | XP_045759629.1 | ENSMPUG00000009122.1 | *PIK3CB* |
| Hfa007 | 39312421 | 39322394 | 23 | XP_045759638.1 | ENSMPUG00000009142.1 | *FAIM* |
| Hfa007 | 39374580 | 39429790 | 23 | XP_045759633.1 | ENSMPUG00000009149.1 | *CEP70* |
| Hfa007 | 39445505 | 39493039 | 23 | XP_045759643.1 | ENSMPUG00000009161.1 | *ESYT3* |
| Hfa007 | 39527376 | 39554404 | 23 | XP_045759644.1 | ENSMPUG00000009182.1 | *MRAS* |
| Hfa007 | 39593264 | 39613322 | 23 | XP_064443476.1 | ENSMPUG00000009189.1 | *NME9* |
| Hfa007 | 39619049 | 39722011 | 23 | XP_045759654.1 | ENSMPUG00000009208.1 | *ARMC8* |
| Hfa007 | 39735549 | 39746148 | 23 | XP_045759658.1 | ENSMPUG00000009230.1 | *DBR1* |
| Hfa007 | 39768943 | 39773344 | 23 | XP_045759660.1 | ENSMPUG00000009238.1 | *A4GNT* |
| Hfa007 | 39793587 | 39824534 | 23 | XP_064443490.1 | ENSMPUG00000009244.1 | *DZIP1L* |
| Hfa007 | 39856676 | 39878153 | 23 | XP_045719676.1 | ENSMPUG00000009256.1 | *CLDN18* |
| Hfa007 | 40035447 | 40036169 | 23 | XP_045719678.1 | ENSMPUG00000018663.1 | *SOX14* |
| Hfa012 | 22608261 | 22914398 | 13 | XP_045751823.1 | ENSMPUG00000001385.1 | *ASAP1* |
| Hfa012 | 23203673 | 23442288 | 13 | XP_064443511.1 | ENSMPUG00000001395.1 | *ADCY8* |
| Hfa012 | 24162476 | 24229905 | 13 | XP_045751820.1 | ENSMPUG00000001401.1 | *EFR3A* |
| Hfa012 | 24236213 | 24265458 | 13 | XP_045751532.1 | ENSMPUG00000001413.1 | *OC90* |
| Hfa012 | 24273340 | 24306301 | 13 | XP_045751530.1 | ENSMPUG00000001421.1 | *HHLA1* |
| Hfa012 | 24328731 | 24375845 | 13 | XP_045751819.2 | ENSMPUG00000001440.1 | *KCNQ3* |
| Hfa012 | 24696205 | 24756356 | 13 | XP_045751817.1 | ENSMPUG00000001571.1 | *DNAAF11* |
| Hfa012 | 24787073 | 24814347 | 13 | XP_045751810.1 | ENSMPUG00000001577.1 | *TMEM71* |
| Hfa020 | 6509170 | 6523439 | 8 | XP_045756609.1 | ENSMPUG00000005358.1 | *COCH* |
| Hfa020 | 6528444 | 6630795 | 8 | XP_045756626.1 | ENSMPUG00000005340.1 | *STRN3* |
| Hfa020 | 6656140 | 6675547 | 8 | XP_045756630.1 | ENSMPUG00000005330.1 | *AP4S1* |
| Hfa020 | 6679482 | 6764122 | 8 | XP_045756610.1 | ENSMPUG00000005315.1 | *HECTD1* |
| Hfa020 | 6837835 | 6930646 | 8 | XP_064426985.1 | ENSMPUG00000001067.1 | *HEATR5A* |
| Hfa020 | 7050268 | 7266321 | 8 | XP_064426998.1 | ENSMPUG00000001061.1 | *NUBPL* |
| Hfa020 | 7469554 | 7532114 | 8 | XP_045756637.1 | ENSMPUG00000001057.1 | *ARHGAP5* |
| Hfa020 | 7779486 | 8142097 | 8 | XP_064427006.1 | ENSMPUG00000001051.1 | *AKAP6* |
| Hfa020 | 8342400 | 9064778 | 8 | XP_064427820.1 | ENSMPUG00000001045.1 | *NPAS3* |
| Hfa023 | 8516155 | 8608668 | 3 | XP_045754086.1 | ENSMPUG00000007964.1 | *MEF2C* |
| Hfa023 | 9008696 | 9053057 | 3 | XP_045753936.1 | ENSMPUG00000007974.1 | *TMEM161B* |
| Hfa027 | 457 | 81401 | 7 | XP_064433849.1 | ENSMPUG00000006957.1 | *TRAPPC8* |
| Hfa027 | 276659 | 283099 | 7 | XP_045742840.1 | ENSMPUG00000006945.1 | *TTR* |
| Hfa027 | 318956 | 364937 | 7 | XP_045742841.2 | ENSMPUG00000006935.1 | *DSG2* |
| Hfa027 | 381222 | 410230 | 7 | XP_045742842.2 | ENSMPUG00000006917.1 | *DSG3* |
| Hfa027 | 434428 | 493356 | 7 | XP_045743907.1 | ENSMPUG00000006860.1 | *DSG4* |
| Hfa027 | 494305 | 516766 | 7 | XP_045742348.2 | ENSMPUG00000006844.1 | *DSG1* |
| Hfa027 | 665309 | 693077 | 7 | XP_064434901.1 | ENSMPUG00000006835.1 | *DSC1* |
| Hfa027 | 713281 | 745687 | 7 | XP_045742349.1 | ENSMPUG00000006720.1 | *DSC2* |
| Hfa027 | 778581 | 811760 | 7 | XP_045743909.3 | ENSMPUG00000006644.1 | *DSC3* |
| Hfa034 | 182485 | 272551 | 8 | XP_045757987.2 | ENSMPUG00000003031.1 | *BCL11B* |
| Hfa040 | 1230233 | 1249099 | 7 | XP_045742287.1 | ENSMPUG00000000750.1 | *LIPG* |

| | | | | | | |
|---|---|---|---|---|---|---|
| Hfa040 | 1333124 | 1650184 | 7 | XP_045742697.1 | ENSMPUG00000000838.1 | *DYM* |
| Hfa040 | 1729568 | 1757409 | 7 | XP_045742703.1 | ENSMPUG00000000842.1 | *SMAD7* |
| Hfa040 | 1808022 | 2016964 | 7 | XP_045742705.1 | ENSMPUG00000000844.1 | *CTIF* |
| Hfa040 | 2525091 | 2536846 | 7 | XP_045742714.1 | ENSMPUG00000000848.1 | *ZBTB7C* |
| Hfa040 | 2643529 | 2695629 | 7 | XP_045742716.1 | ENSMPUG00000000850.1 | *SMAD2* |
| Hfa040 | 3171387 | 3204891 | 7 | XP_045742288.1 | ENSMPUG00000000851.1 | |
| Hfa045 | 1902782 | 1904353 | 4 | XP_045732283.1 | ENSMPUG00000011947.1 | *LRRTM3* |
| Hfa049 | 5510327 | 6311836 | X | XP_064429968.1 | ENSMPUG00000002354.1 | *DMD* |
| Hfa055 | 3676704 | 4459316 | 15 | XP_064442473.1 | ENSMPUG00000001059.1 | *CNTNAP5* |
| Hfa058 | 13733 | 29562 | 11 | XP_045738660.1 | ENSMPUG00000006040.1 | *POLR1E* |
| Hfa058 | 46927 | 94552 | 11 | XP_064435254.1 | ENSMPUG00000006111.1 | *FBXO10* |
| Hfa058 | 210286 | 261022 | 11 | XP_045738666.1 | ENSMPUG00000006120.1 | *FRMPD1* |
| Hfa058 | 265152 | 280682 | 11 | XP_064435282.1 | ENSMPUG00000006133.1 | *TRMT10B* |
| Hfa058 | 282542 | 288203 | 11 | XP_045738670.1 | ENSMPUG00000006144.1 | *EXOSC3* |
| Hfa058 | 312336 | 350867 | 11 | XP_045738671.1 | ENSMPUG00000006153.1 | *DCAF10* |
| Hfa058 | 397024 | 529390 | 11 | XP_045738683.1 | ENSMPUG00000006187.1 | *SHB* |
| Hfa058 | 758124 | 759676 | 11 | XP_054365289.2 | ENSMPUG00000018536.1 | *ALDH1B1* |
| Hfa058 | 794956 | 841058 | 11 | XP_045738684.1 | ENSMPUG00000006197.1 | |
| Hfa058 | 851209 | 913406 | 11 | XP_064436104.1 | ENSMPUG00000006330.1 | *CCDC180* |
| Hfa058 | 942761 | 1003094 | 11 | XP_045738694.1 | ENSMPUG00000006366.1 | *TDRD7* |
| Hfa058 | 1027936 | 1089292 | 11 | XP_045738709.1 | ENSMPUG00000006375.1 | *TMOD1* |
| Hfa058 | 1086747 | 1110937 | 11 | XP_045738707.2 | ENSMPUG00000006386.1 | *TSTD2* |
| Hfa066 | 17632 | 73357 | 6 | XP_064449419.1 | ENSMPUG00000016613.1 | *FBXL18* |
| Hfa066 | 90461 | 173454 | 6 | XP_045731836.2 | ENSMPUG00000016618.1 | *TNRC18* |
| Hfa066 | 177079 | 187074 | 6 | XP_045728164.1 | ENSMPUG00000016628.1 | *SLC29A4* |
| Hfa066 | 224520 | 260187 | 6 | XP_045731839.1 | ENSMPUG00000016629.1 | *WIPI2* |
| Hfa066 | 276878 | 316927 | 6 | XP_045726969.1 | ENSMPUG00000016644.1 | *MMD2* |
| Hfa066 | 336293 | 393676 | 6 | XP_045731841.1 | ENSMPUG00000016653.1 | *RADIL* |
| Hfa066 | 396629 | 408694 | 6 | XP_064449432.1 | ENSMPUG00000016654.1 | *AP5Z1* |
| Hfa066 | 417207 | 477206 | 6 | XP_045731846.1 | ENSMPUG00000016655.1 | *FOXK1* |
| Hfa066 | 669534 | 1196248 | 6 | XP_054363696.1 | ENSMPUG00000016658.1 | |
| Hfa082 | 8730833 | 8745743 | X | XP_045740834.1 | ENSMPUG00000001193.1 | *ASMT* |
| Hfa082 | 8757222 | 8764658 | X | XP_045729861.2 | ENSMPUG00000001208.1 | *AKAP17A* |
| Hfa082 | 8841811 | 8864525 | X | XP_045729850.2 | ENSMPUG00000001209.1 | *ASMTL* |
| Hfa082 | 8876160 | 8894880 | X | XP_064429734.1 | ENSMPUG00000001223.1 | |
| Hfa082 | 8943411 | 8963519 | X | XP_045729791.2 | ENSMPUG00000001235.1 | *CSF2RA* |
| Hfa082 | 8998841 | 9020860 | X | XP_045728539.1 | ENSMPUG00000001250.1 | *CRLF2* |
| Hfa082 | 9393228 | 9402817 | X | XP_045740504.1 | ENSMPUG00000001254.1 | *SHOX* |
| Hfa082 | 9754097 | 9765928 | X | XP_064429728.1 | ENSMPUG00000001536.1 | *GTPBP6* |
| Hfa098 | 6350005 | 6368573 | 6 | XP_045753386.1 | ENSMPUG00000000887.1 | *DEPDC1* |
| Hfa098 | 6401934 | 6421366 | 6 | XP_045751356.1 | ENSMPUG00000000810.1 | *RPE65* |
| Hfa098 | 6574226 | 6671408 | 6 | XP_045753389.1 | ENSMPUG00000000807.1 | *WLS* |
| Hfa098 | 7013157 | 7014929 | 6 | XP_045753396.1 | ENSMPUG00000000798.1 | *GADD45A* |
| Hfa125 | 4014249 | 4015085 | 2 | XP_045750602.1 | ENSMPUG00000020044.1 | *KCTD16* |
| Hfa125 | 4050028 | 4061556 | 2 | XP_045750566.1 | ENSMPUG00000007876.1 | *YIPF5* |
| Hfa142 | 687353 | 687979 | X | XP_045732480.1 | ENSMPUG00000018947.1 | *PPP1R2C* |
| Hfa142 | 1613206 | 1623346 | X | XP_045746066.1 | ENSMPUG00000012912.1 | *NDP* |
| Hfa142 | 1801832 | 1992997 | X | XP_045732544.1 | ENSMPUG00000012914.1 | *EFHC2* |
| Hfa142 | 2137322 | 2149601 | X | XP_064430017.1 | ENSMPUG00000012924.1 | *FUNDC1* |
| Hfa142 | 2367686 | 2567961 | X | XP_064430018.1 | ENSMPUG00000012926.1 | *KDM6A* |
| Hfa171 | 780672 | 949273 | 1 | XP_064435932.1 | ENSMPUG00000005120.1 | |
| Hfa171 | 1095114 | 1250656 | 1 | XP_054365125.1 | ENSMPUG00000005101.1 | *RFX3* |
| Hfa171 | 2233865 | 2307298 | 1 | XP_045738133.1 | ENSMPUG00000005081.1 | *SLC1A1* |
| Hfa171 | 2328643 | 2370135 | 1 | XP_064436064.1 | ENSMPUG00000005024.1 | *SPATA6L* |
| Hfa171 | 2382869 | 2408051 | 1 | XP_045738128.1 | ENSMPUG00000005018.1 | *CDC37L1* |
| Hfa171 | 2464529 | 2525320 | 1 | XP_045738121.1 | ENSMPUG00000004998.1 | *RCL1* |
| Hfa171 | 2720111 | 2797748 | 1 | XP_045738116.1 | ENSMPUG00000004974.1 | *JAK2* |
| Hfa178 | 856317 | 1112418 | X | XP_064430750.1 | ENSMPUG00000011725.1 | *GABRA3* |
| Hfa178 | 1208758 | 1224620 | X | XP_045739724.1 | ENSMPUG00000011717.1 | *GABRE* |
| Hfa178 | 1462680 | 1468755 | X | XP_045739685.1 | ENSMPUG00000011689.1 | *CNGA2* |
| Hfa178 | 1480268 | 1487548 | X | XP_045739708.1 | ENSMPUG00000011674.1 | *FATE1* |
| Hfa178 | 1505387 | 1506453 | X | XP_045739688.1 | ENSMPUG00000011673.1 | *PRRG3* |
| Hfa178 | 1860134 | 1864797 | X | XP_045727338.2 | ENSMPUG00000011657.1 | *GPR50* |
| Hfa178 | 2038244 | 2122226 | X | XP_045739647.2 | ENSMPUG00000011599.1 | *CD99L2* |
| Hfa178 | 2127032 | 2171233 | X | XP_054365463.1 | ENSMPUG00000011593.1 | *MTMR1* |
| Hfa178 | 2203733 | 2273554 | X | XP_045739602.1 | ENSMPUG00000011585.1 | *MTM1* |
| Hfa178 | 2348805 | 2403368 | X | XP_064430739.1 | ENSMPUG00000011578.1 | *MAMLD1* |
| Hfa209 | 775596 | 799075 | X | XP_045734568.1 | ENSMPUG00000011421.1 | *GNL3L* |
| Hfa209 | 828188 | 868066 | X | XP_045734537.1 | ENSMPUG00000011428.1 | *FGD1* |
| Hfa209 | 869583 | 873648 | X | XP_045734547.1 | ENSMPUG00000011436.1 | *TSR2* |
| Hfa209 | 966895 | 1110498 | X | XP_064430174.1 | ENSMPUG00000011455.1 | *WNK3* |
| Hfa209 | 1273260 | 1359466 | X | XP_045734417.1 | ENSMPUG00000011546.1 | *PHF8* |

| | | | | | | |
|---|---|---|---|---|---|---|
| Hfa209 | 1625436 | 1756916 | X | XP_045734349.2 | ENSMPUG00000009760.1 | *HUWE1* |
| Hfa209 | 1835086 | 1837269 | X | XP_045734326.1 | ENSMPUG00000009755.1 | *HSD17B10* |
| Hfa209 | 1837712 | 1846456 | X | XP_045734306.1 | ENSMPUG00000009750.1 | *RIBC1* |
| Hfa209 | 1849841 | 1882143 | X | XP_045734252.1 | ENSMPUG00000009735.1 | *SMC1A* |
| Hfa209 | 1936035 | 2010655 | X | XP_045734239.1 | ENSMPUG00000009723.1 | *IQSEC2* |
| Hfa209 | 2113462 | 2119979 | X | XP_045734148.1 | ENSMPUG00000009638.1 | *TSPYL2* |
| Hfa220 | 546247 | 552295 | 1 | XP_045746356.2 | ENSMPUG00000006390.1 | *CEACAM16* |
| Hfa220 | 589895 | 604172 | 1 | XP_045747667.1 | ENSMPUG00000006405.1 | *PVR* |
| Hfa220 | 613881 | 628742 | 1 | XP_045748131.2 | ENSMPUG00000006436.1 | |
| Hfa220 | 724680 | 739284 | 1 | XP_054367429.1 | ENSMPUG00000006445.1 | *ZNF180* |
| Hfa220 | 771204 | 780947 | 1 | XP_045747662.2 | ENSMPUG00000006454.1 | *ZNF229* |
| Hfa220 | 792529 | 803738 | 1 | XP_064436518.1 | ENSMPUG00000006463.1 | *ZNF285* |
| Hfa220 | 844596 | 864593 | 1 | XP_045747636.2 | ENSMPUG00000006469.1 | *ZNF112* |
| Hfa220 | 1006076 | 1025309 | 1 | XP_045747639.1 | ENSMPUG00000006485.1 | *ZNF227* |
| Hfa220 | 1042661 | 1052366 | 1 | XP_064437366.1 | ENSMPUG00000006493.1 | *ZNF226* |
| Hfa220 | 1064521 | 1074171 | 1 | XP_045747655.1 | ENSMPUG00000006501.1 | *ZNF234* |
| Hfa220 | 1092808 | 1105484 | 1 | XP_064437585.1 | ENSMPUG00000006510.1 | |
| Hfa220 | 1130755 | 1140825 | 1 | XP_054367532.2 | ENSMPUG00000006522.1 | *ZNF45* |
| Hfa220 | 1186533 | 1193864 | 1 | XP_045747656.1 | ENSMPUG00000006528.1 | *ZNF404* |
| Hfa220 | 1289574 | 1293381 | 1 | XP_045747634.1 | ENSMPUG00000006560.1 | *LYPD5* |
| Hfa220 | 1305643 | 1317134 | 1 | XP_045747632.1 | ENSMPUG00000006586.1 | *KCNN4* |
| Hfa220 | 1329760 | 1347011 | 1 | XP_045747628.1 | ENSMPUG00000006594.1 | *SMG9* |
| Hfa220 | 1354196 | 1355590 | 1 | XP_064436523.1 | ENSMPUG00000006607.1 | *IRGC* |
| Hfa220 | 1390876 | 1403837 | 1 | XP_045747627.1 | ENSMPUG00000006628.1 | *PLAUR* |
| Hfa220 | 1411924 | 1424713 | 1 | XP_045746353.1 | ENSMPUG00000006662.1 | *CADM4* |
| Hfa220 | 1430606 | 1436171 | 1 | XP_045747624.1 | ENSMPUG00000006718.1 | *ZNF428* |
| Hfa220 | 1443021 | 1446066 | 1 | XP_045747621.1 | ENSMPUG00000006726.1 | *ZNF576* |
| Hfa220 | 1446454 | 1449728 | 1 | XP_045747595.1 | ENSMPUG00000006734.1 | *IRGQ* |
| Hfa220 | 1456366 | 1460094 | 1 | XP_045747606.1 | ENSMPUG00000006741.1 | *PINLYP* |
| Hfa220 | 1461374 | 1486971 | 1 | XP_045747601.1 | ENSMPUG00000006750.1 | *XRCC1* |
| Hfa220 | 1498430 | 1515942 | 1 | XP_045747608.1 | ENSMPUG00000006865.1 | *ETHE1* |
| Hfa220 | 1518002 | 1536574 | 1 | XP_045747594.1 | ENSMPUG00000006905.1 | *PHLDB3* |
| Hfa220 | 1542060 | 1545655 | 1 | XP_045748126.1 | ENSMPUG00000006923.1 | *LYPD3* |
| Hfa220 | 1592054 | 1605907 | 1 | XP_064437694.1 | ENSMPUG00000006959.1 | *CD177* |
| Hfa220 | 1636758 | 1646239 | 1 | XP_064438107.1 | ENSMPUG00000007009.1 | |
| Hfa220 | 1703352 | 1704999 | 1 | XP_045746351.1 | ENSMPUG00000007126.1 | *LYPD4* |
| Hfa220 | 1710662 | 1714358 | 1 | XP_045746349.1 | ENSMPUG00000007147.1 | *DMRTC2* |
| Hfa220 | 1721809 | 1729849 | 1 | XP_045747589.1 | ENSMPUG00000007164.1 | *RPS19* |
| Hfa220 | 1732885 | 1736656 | 1 | XP_045746348.2 | ENSMPUG00000007182.1 | *CD79A* |
| Hfa220 | 1745689 | 1758894 | 1 | XP_045747586.1 | ENSMPUG00000007235.1 | *ARHGEF1* |
| Hfa220 | 1760261 | 1764902 | 1 | XP_045747588.1 | ENSMPUG00000007268.1 | *ERFL* |
| Hfa220 | 1800188 | 1802620 | 1 | XP_045747580.1 | ENSMPUG00000007273.1 | *RABAC1* |
| Hfa220 | 1807879 | 1822356 | 1 | XP_045748121.2 | ENSMPUG00000007324.1 | *ATP1A3* |
| Hfa220 | 1831863 | 1880838 | 1 | XP_045747578.1 | ENSMPUG00000007345.1 | *GRIK5* |
| Hfa221 | 528 | 7134 | 6 | XP_045731427.1 | ENSMPUG00000016192.1 | *SBDS* |
| Hfa221 | 60904 | 71911 | 6 | XP_064449142.1 | ENSMPUG00000016193.1 | *TMEM248* |
| Hfa221 | 117759 | 200944 | 6 | XP_064449111.1 | ENSMPUG00000016195.1 | *RABGEF1* |
| Hfa221 | 127366 | 200944 | 6 | XP_064449116.1 | ENSMPUG00000016196.1 | *KCTD7* |
| Hfa221 | 214879 | 327535 | 6 | XP_064449125.1 | ENSMPUG00000016197.1 | *TPST1* |
| Hfa221 | 437413 | 477579 | 6 | XP_064449138.1 | ENSMPUG00000016198.1 | *CRCP* |
| Hfa221 | 511578 | 517594 | 6 | XP_054362536.2 | ENSMPUG00000016200.1 | *ASL* |
| Hfa221 | 542508 | 555405 | 6 | XP_045731409.2 | ENSMPUG00000016201.1 | *GUSB* |
| Hfa221 | 722145 | 727857 | 6 | XP_045731402.1 | ENSMPUG00000016208.1 | *PHKG1* |
| Hfa221 | 729022 | 739311 | 6 | XP_054363590.2 | ENSMPUG00000016209.1 | *SUMF2* |
| Hfa221 | 793837 | 800430 | 6 | XP_045731393.1 | ENSMPUG00000016227.1 | *PSPH* |
| Hfa221 | 808427 | 837625 | 6 | XP_045731389.1 | ENSMPUG00000016229.1 | *NIPSNAP2* |
| Hfa221 | 840942 | 843121 | 6 | XP_045731379.1 | ENSMPUG00000016230.1 | *MRPS17* |
| Hfa221 | 887605 | 926155 | 6 | XP_064449100.1 | ENSMPUG00000016234.1 | *ZNF713* |
| Hfa221 | 971718 | 1017268 | 6 | XP_064446450.1 | ENSMPUG00000010720.1 | *SEPTIN10* |
| Hfa221 | 1064737 | 1065237 | 6 | XP_045727943.1 | ENSMPUG00000011332.1 | *AHSP* |
| Hfa221 | 1073358 | 1089583 | 6 | XP_045731373.1 | ENSMPUG00000011335.1 | *RUSF1* |
| Hfa221 | 1088455 | 1094436 | 6 | XP_045731376.2 | ENSMPUG00000011343.1 | *SLC5A2* |
| Hfa221 | 1098209 | 1101868 | 6 | XP_045731374.1 | ENSMPUG00000011350.1 | *TGFB1I1* |
| Hfa221 | 1105131 | 1110776 | 6 | XP_045731371.1 | ENSMPUG00000011355.1 | *ARMC5* |
| Hfa221 | 1141181 | 1141774 | 6 | XP_045731369.2 | ENSMPUG00000011359.1 | *COX6A2* |
| Hfa221 | 1292611 | 1345679 | 6 | XP_045731357.1 | ENSMPUG00000011385.1 | *ITGAM* |
| Hfa221 | 1408890 | 1409867 | 6 | XP_045738395.1 | ENSMPUG00000004058.1 | *PSIP1* |
| Hfa221 | 1413089 | 1419791 | 6 | XP_045731356.2 | ENSMPUG00000011397.1 | *TRIM72* |
| Hfa221 | 1429951 | 1431338 | 6 | XP_045727917.1 | ENSMPUG00000011423.1 | *PYCARD* |
| Hfa221 | 1440415 | 1449258 | 6 | XP_045731352.1 | ENSMPUG00000011426.1 | |
| Hfa221 | 1475370 | 1486712 | 6 | XP_045727905.1 | ENSMPUG00000011434.1 | *PRSS36* |
| Hfa221 | 1493568 | 1502971 | 6 | XP_045731351.1 | ENSMPUG00000011488.1 | *KAT8* |

| | | | | | | |
|---|---|---|---|---|---|---|
| Hfa221 | 1506925 | 1509960 | 6 | XP_045731345.1 | ENSMPUG00000011540.1 | *BCKDK* |
| Hfa221 | 1521574 | 1526377 | 6 | XP_045731350.2 | ENSMPUG00000011621.1 | *PRSS53* |
| Hfa221 | 1522609 | 1533931 | 6 | XP_064449075.1 | ENSMPUG00000011652.1 | *ZNF646* |
| Hfa221 | 1541760 | 1544186 | 6 | XP_045731338.1 | ENSMPUG00000011656.1 | *ZNF668* |
| Hfa221 | 1561224 | 1565723 | 6 | XP_045731337.1 | ENSMPUG00000011670.1 | *STX4* |
| Hfa221 | 1577977 | 1592901 | 6 | XP_045731336.2 | ENSMPUG00000011677.1 | *STX1B* |
| Hfa221 | 1597066 | 1599946 | 6 | XP_045731334.1 | ENSMPUG00000011815.1 | *HSD3B7* |
| Hfa221 | 1600972 | 1621217 | 6 | XP_045731329.1 | ENSMPUG00000011867.1 | |
| Hfa221 | 1625144 | 1629210 | 6 | XP_045731335.1 | ENSMPUG00000011954.1 | *ORAI3* |
| Hfa221 | 1631429 | 1647699 | 6 | XP_045731332.1 | ENSMPUG00000011968.1 | *FBXL19* |
| Hfa221 | 1668520 | 1670773 | 6 | XP_064446395.1 | ENSMPUG00000011977.1 | |
| Hfa221 | 1672244 | 1675711 | 6 | XP_045731326.1 | ENSMPUG00000011973.1 | |
| Hfa221 | 1678319 | 1717993 | 6 | XP_045731324.1 | ENSMPUG00000011988.1 | *BCL7C* |
| Hfa221 | 1776999 | 1787870 | 6 | XP_045731310.1 | ENSMPUG00000012016.1 | *RNF40* |
| Hfa221 | 1788614 | 1792152 | 6 | XP_045731318.1 | ENSMPUG00000012128.1 | *CFAP119* |
| Hfa221 | 1792535 | 1801553 | 6 | XP_045731316.1 | ENSMPUG00000012139.1 | *PHKG2* |
| Hfa226 | 320510 | 543660 | 9 | XP_064439795.1 | ENSMPUG00000012831.1 | *RAP1GAP2* |
| Hfa226 | 819273 | 820436 | 9 | XP_064438194.1 | ENSMPUG00000018512.1 | |
| Hfa226 | 900602 | 916963 | 9 | XP_064439733.1 | ENSMPUG00000006045.1 | *ASPA* |
| Hfa226 | 924678 | 956222 | 9 | XP_045744102.2 | ENSMPUG00000005865.1 | |
| Hfa226 | 964569 | 991446 | 9 | XP_045745621.1 | ENSMPUG00000012097.1 | *TRPV1* |
| Hfa226 | 1004268 | 1027650 | 9 | XP_045745615.2 | ENSMPUG00000012193.1 | *SHPK* |
| Hfa226 | 1030695 | 1044342 | 9 | XP_064438386.1 | ENSMPUG00000012199.1 | *CTNS* |
| Hfa226 | 1047399 | 1051425 | 9 | XP_064438388.1 | ENSMPUG00000012203.1 | *TAX1BP3* |
| Hfa226 | 1051989 | 1052321 | 9 | XP_045745622.1 | ENSMPUG00000018553.1 | *EMC6* |
| Hfa226 | 1058080 | 1069758 | 9 | XP_045745623.1 | ENSMPUG00000012236.1 | *P2RX5* |
| Hfa226 | 1088587 | 1158120 | 9 | XP_045745629.2 | ENSMPUG00000012243.1 | *ITGAE* |
| Hfa226 | 1158138 | 1198707 | 9 | XP_064438788.1 | ENSMPUG00000012357.1 | *NCBP3* |
| Hfa226 | 1209828 | 1228065 | 9 | XP_045745630.1 | ENSMPUG00000012363.1 | *CAMKK1* |
| Hfa226 | 1239809 | 1254218 | 9 | XP_064439495.1 | ENSMPUG00000012367.1 | *P2RX1* |
| Hfa226 | 1261409 | 1294722 | 9 | XP_045745633.1 | ENSMPUG00000012441.1 | *ATP2A3* |
| Hfa226 | 1650556 | 1652311 | 5 | XP_064438962.1 | ENSMPUG00000019373.1 | *KCNJ12* |
| Hfa226 | 1749608 | 1762690 | 5 | XP_045745639.1 | ENSMPUG00000000007.1 | *MAP2K3* |
| Hfa231 | 54433 | 71696 | 2 | XP_045754296.1 | ENSMPUG00000015601.1 | *WASF2* |
| Hfa231 | 84034 | 85026 | 2 | XP_045751430.1 | ENSMPUG00000018813.1 | *GPR3* |
| Hfa231 | 92377 | 95542 | 2 | XP_045751431.1 | ENSMPUG00000015613.1 | *CD164L2* |
| Hfa231 | 104808 | 116803 | 2 | XP_064443624.1 | ENSMPUG00000015615.1 | *MAP3K6* |
| Hfa231 | 117288 | 123675 | 2 | XP_064445771.1 | ENSMPUG00000015617.1 | *SYTL1* |
| Hfa231 | 129415 | 140061 | 2 | XP_045754300.1 | ENSMPUG00000015642.1 | *TMEM222* |
| Hfa231 | 151354 | 195665 | 2 | XP_045754302.1 | ENSMPUG00000015651.1 | *WDTC1* |
| Hfa231 | 264962 | 311740 | 2 | XP_064445775.1 | ENSMPUG00000015654.1 | *SLC9A1* |
| Hfa231 | 371868 | 378056 | 2 | XP_045754307.1 | ENSMPUG00000015656.1 | *TENT5B* |
| Hfa231 | 419916 | 422160 | 2 | XP_045751433.1 | ENSMPUG00000015660.1 | *KDF1* |
| Hfa231 | 424166 | 437899 | 2 | XP_045754309.1 | ENSMPUG00000015663.1 | *NUDC* |
| Hfa231 | 448729 | 450352 | 2 | XP_045751434.1 | ENSMPUG00000015664.1 | *NR0B2* |
| Hfa231 | 456103 | 462879 | 2 | XP_045754310.1 | ENSMPUG00000015665.1 | *GPATCH3* |
| Hfa231 | 463786 | 471453 | 2 | XP_045754311.1 | ENSMPUG00000015666.1 | *GPN2* |
| Hfa231 | 482008 | 482754 | 2 | XP_045754313.2 | ENSMPUG00000018816.1 | *SFN* |
| Hfa231 | 490987 | 512198 | 2 | XP_045754312.1 | ENSMPUG00000015672.1 | *ZDHHC18* |
| Hfa231 | 525598 | 531250 | 2 | XP_045754314.2 | ENSMPUG00000015674.1 | *PIGV* |
| Hfa231 | 542099 | 608335 | 2 | XP_045754319.1 | ENSMPUG00000015675.1 | *ARID1A* |
| Hfa231 | 779514 | 782401 | 2 | XP_045726286.1 | ENSMPUG00000008613.1 | *HMGN4* |
| Hfa231 | 785797 | 813129 | 2 | XP_045726278.1 | ENSMPUG00000015707.1 | *DHDDS* |
| Hfa231 | 818257 | 830288 | 2 | XP_045726276.1 | ENSMPUG00000015724.1 | *LIN28A* |
| Hfa231 | 869608 | 895873 | 2 | XP_045754327.2 | ENSMPUG00000015728.1 | *CRYBG2* |
| Hfa231 | 915830 | 933463 | 2 | XP_045754329.1 | ENSMPUG00000015736.1 | *UBXN11* |
| Hfa231 | 934542 | 935597 | 2 | XP_045754333.1 | ENSMPUG00000015740.1 | *SH3BGRL3* |
| Hfa231 | 938424 | 963169 | 2 | XP_045754334.1 | ENSMPUG00000015744.1 | *CEP85* |
| Hfa231 | 982517 | 995409 | 2 | XP_054369206.1 | ENSMPUG00000015760.1 | *CATSPER4* |
| Hfa231 | 996065 | 1005184 | 2 | XP_045754338.1 | ENSMPUG00000015762.1 | *CNKSR1* |
| Hfa231 | 1018966 | 1023972 | 2 | XP_064445802.1 | ENSMPUG00000015764.1 | *FAM110D* |
| Hfa231 | 1040738 | 1047004 | 2 | XP_045754343.1 | ENSMPUG00000015769.1 | *PDIK1L* |
| Hfa231 | 1078738 | 1088736 | 2 | XP_045754346.1 | ENSMPUG00000015770.1 | *TRIM63* |
| Hfa231 | 1103487 | 1109787 | 2 | XP_045751439.1 | ENSMPUG00000015771.1 | *SLC30A2* |
| Hfa231 | 1113214 | 1126183 | 2 | XP_045754347.1 | ENSMPUG00000015772.1 | *EXTL1* |
| Hfa231 | 1150735 | 1173032 | 2 | XP_045754348.1 | ENSMPUG00000015773.1 | *PAFAH2* |
| Hfa231 | 1268648 | 1269694 | 2 | XP_045754353.1 | ENSMPUG00000018819.1 | *PAQR7* |
| Hfa231 | 1272819 | 1294745 | 2 | XP_045754356.2 | ENSMPUG00000015775.1 | *AUNIP* |
| Hfa231 | 1296439 | 1308997 | 2 | XP_045754354.1 | ENSMPUG00000015776.1 | *MTFR1L* |
| Hfa231 | 1344596 | 1480650 | 2 | XP_045754359.1 | ENSMPUG00000015777.1 | *MAN1C1* |
| Hfa231 | 1518014 | 1539682 | 2 | XP_045754362.1 | ENSMPUG00000015779.1 | *LDLRAP1* |
| Hfa231 | 1573126 | 1630427 | 2 | XP_045754365.1 | ENSMPUG00000015792.1 | *MACO1* |

| | | | | | | |
|---|---|---|---|---|---|---|
| Hfa411 | 18100 | 52220 | X | XP_064449807.1 | ENSMPUG00000004216.1 | *IQGAP3* |
| Hfa411 | 87857 | 89595 | X | XP_045731997.1 | ENSMPUG00000004131.1 | *HAPLN2* |
| Hfa411 | 105169 | 117676 | X | XP_045731998.1 | ENSMPUG00000004123.1 | *BCAN* |
| Hfa411 | 125768 | 134209 | X | XP_045732606.1 | ENSMPUG00000004103.1 | *NES* |
| Hfa411 | 166001 | 169521 | X | XP_045732616.1 | ENSMPUG00000004066.1 | *ISG20L2* |

Table S3. Genes within 1 Mbp of regions or variants with $\Delta AF \geq 0.55$ and Gene Ontology (GO) terms related mainly to ion homeostasis and lipid metabolism. Only a subset of GO names is shown for each gene.

| Contig | Start | End | Gene symbol | GO ID (subset) | GO names |
|---|---|---|---|---|---|
| Hfa001 | 46045770 | 46072559 | *PI4K2B* | GO:0006629 | lipid metabolic process |
| Hfa007 | 38439798 | 38463659 | *RBP1* | GO:0005811; GO:0019915; GO:0008289 | lipid droplet; lipid storage; lipid binding |
| Hfa007 | 38503646 | 38525870 | *RBP2* | GO:0008289 | lipid binding |
| Hfa007 | 39186568 | 39297343 | *PIK3CB* | GO:0006629 | lipid metabolic process |
| Hfa007 | 39445505 | 39493039 | *ESYT3* | GO:0005544; GO:0008289; GO:0006869 | calcium-dependent phospholipid binding; lipid binding; lipid transport |
| Hfa012 | 24328731 | 24375845 | *KCNQ3* | GO:0008076; GO:0071805; GO:0034702 | voltage-gated potassium channel complex; potassium ion transmembrane transport; monoatomic ion channel complex |
| Hfa020 | 7779486 | 8142097 | *AKAP6* | GO:0010959; GO:1904064; GO:1901381 | regulation of metal ion transport; positive regulation of cation transmembrane transport; positive regulation of potassium ion transmembrane transport |
| Hfa034 | 182485 | 272551 | *BCL11B* | GO:0019216 | regulation of lipid metabolic process |
| Hfa040 | 1230233 | 1249099 | *LIPG* | GO:0004465; GO:0016042; GO:0006629 | lipoprotein lipase activity; lipid catabolic process; lipid metabolic process |
| Hfa049 | 5510327 | 6311836 | *DMD* | GO:1902305 | regulation of sodium ion transmembrane transport |
| Hfa066 | 177079 | 187074 | *SLC29A4* | GO:0098655; GO:0015695; GO:0015101 | monoatomic cation transmembrane transport; organic cation transport; organic cation transmembrane transporter activity |
| Hfa098 | 7013157 | 7014929 | *GADD45A* | GO:0051726; GO:1900745; GO:0046330 | regulation of cell cycle; positive regulation of p38MAPK cascade; positive regulation of JNK cascade |
| Hfa171 | 2233865 | 2307298 | *SLC1A1* | GO:1902476; GO:0015501; GO:0015108 | chloride transmembrane transport; glutamate:sodium symporter activity; chloride transmembrane transporter activity |
| Hfa178 | 856317 | 1112418 | *GABRA3* | GO:0022851; GO:0006811; GO:1902476 | GABA-gated chloride ion channel activity; monoatomic ion transport; chloride transmembrane transport |
| Hfa178 | 1208758 | 1224620 | *GABRE* | GO:0022851; GO:2001226; GO:0006811 | GABA-gated chloride ion channel activity; negative regulation of chloride transport; monoatomic ion transport |
| Hfa178 | 1462680 | 1468755 | *CNGA2* | GO:0098655; GO:0005223; GO:0006816 | monoatomic cation transmembrane transport; intracellularly cGMP-activated cation channel activity; calcium ion transport |
| Hfa178 | 2127032 | 2171233 | *MTMR1* | GO:0006629 | lipid metabolic process |
| Hfa178 | 2203733 | 2273554 | *MTM1* | GO:0006629 | lipid metabolic process |
| Hfa209 | 966895 | 1110498 | *WNK3* | GO:0007231; GO:0010765; GO:0070294 | osmosensory signaling pathway; positive regulation of sodium ion transport; renal sodium ion absorption |
| Hfa220 | 1305643 | 1317134 | *KCNN4* | GO:0006813; GO:0015269; GO:0034220 | potassium ion transport; calcium-activated potassium channel activity; monoatomic ion transmembrane transport |
| Hfa220 | 1807879 | 1822356 | *ATP1A3* | GO:0030007; GO:0006883; GO:0005391; | intracellular potassium ion homeostasis; intracellular sodium ion homeostasis; P-type sodium:potassium-exchanging transporter activity |
| Hfa220 | 1831863 | 1880838 | *GRIK5* | GO:0015276; GO:0006811; GO:0005216 | ligand-gated monoatomic ion channel activity; monoatomic ion transport; monoatomic ion channel activity |
| Hfa221 | 127366 | 200944 | *KCTD7* | GO:0030007 | intracellular potassium ion homeostasis |
| Hfa221 | 1088455 | 1094436 | *SLC5A2* | GO:0006814; GO:0005412; GO:0006811 | sodium ion transport; D-glucose:sodium symporter activity; monoatomic ion transport |
| Hfa221 | 1506925 | 1509960 | *BCKDK* | GO:0008610 | lipid biosynthetic process |
| Hfa221 | 1597066 | 1599946 | *HSD3B7* | GO:0005811; GO:0006629 | lipid droplet; lipid metabolic process |
| Hfa226 | 924678 | 956222 | | GO:0007231; GO:0006811; GO:0005216 | osmosensory signaling pathway; monoatomic ion transport; monoatomic ion channel activity |
| Hfa226 | 964569 | 991446 | *TRPV1* | GO:0005230; GO:0006811; GO:0005216 | extracellular ligand-gated monoatomic ion channel activity; monoatomic ion transport; monoatomic ion channel activity |
| Hfa226 | 1058080 | 1069758 | *P2RX5* | GO:0006821; GO:0099095; GO:0035381 | chloride transport; ligand-gated monoatomic anion channel activity; ATP-gated ion channel activity |
| Hfa226 | 1239809 | 1254218 | *P2RX1* | GO:0051924; GO:0005261; GO:0004931 | regulation of calcium ion transport; monoatomic cation channel activity; extracellularly ATP-gated monoatomic cation channel activity |
| Hfa226 | 1261409 | 1294722 | *ATP2A3* | GO:1903515; GO:0006811; GO:0006816 | calcium ion transport from cytosol to endoplasmic reticulum; monoatomic ion transport; calcium ion transport |
| Hfa226 | 1650556 | 1652311 | *KCNJ12* | GO:0005242; GO:0034765; GO:1990573 | inward rectifier potassium channel activity; regulation of monoatomic ion transmembrane transport; potassium ion import across plasma membrane |
| Hfa231 | 84034 | 85026 | *GPR3* | GO:0001659; GO:1990845; GO:0001659 | G protein-coupled receptor activity; adaptive thermogenesis; temperature homeostasis |
| Hfa231 | 264962 | 311740 | *SLC9A1* | GO:0006883; GO:0015385; GO:0015386 | intracellular sodium ion homeostasis; sodium:proton antiporter activity; potassium:proton antiporter activity |
| Hfa231 | 785797 | 813129 | *DHDDS* | GO:0006629 | lipid metabolic process |
| Hfa231 | 982517 | 995409 | *CATSPER4* | GO:0006814; GO:0006811; GO:0005227 | sodium ion transport; monoatomic ion transport; calcium-activated cation channel activity |
| Hfa231 | 1103487 | 1109787 | *SLC30A2* | GO:0006811; GO:0008324; GO:0006829 | monoatomic ion transport; monoatomic cation transport; zinc ion transport |
| Hfa231 | 1150735 | 1173032 | *PAFAH2* | GO:0006629; GO:0016042 | lipid metabolic process; lipid catabolic process |
| Hfa231 | 1518014 | 1539682 | *LDLRAP1* | GO:1905581; GO:0090118 | positive regulation of low-density lipoprotein particle clearance; receptor-mediated endocytosis involved in cholesterol transport |

Table S4. Studied grey seal individuals, with subspecies, sampling location and country, site code, sex, age, type (live, hunted, stranded), sampling/acquisition year, and sequencing depth information. The latter corresponds to the average depth of 50 kbp windows across the largest contig (Hfa001) for each individual.

| Individual | Subspecies | Location | Country | Site code | Sex | Age | Type | Year | Depth |
|---|---|---|---|---|---|---|---|---|---|
| HG543 | Atlantic | Breiðafjörður | Iceland (NW) | ICE_BRE | m | N/A | stranded | 2009 | 15.73 |
| HG544 | Atlantic | Breiðafjörður | Iceland (NW) | ICE_BRE | f | N/A | stranded | 2008 | 15.37 |
| HG545 | Atlantic | Breiðafjörður | Iceland (NW) | ICE_BRE | f | N/A | stranded | 2008 | 17.89 |
| HG546 | Atlantic | Breiðafjörður | Iceland (NW) | ICE_BRE | m | N/A | stranded | 2009 | 12.88 |
| HG550 | Atlantic | Djúpavogur | Iceland (NE) | ICE_DJU | f | N/A | stranded | 2009 | 24.24 |
| HG549 | Atlantic | Djúpivogur | Iceland (NE) | ICE_DJU | f | N/A | stranded | 2009 | 14.08 |
| HG552 | Atlantic | Djúpivogur | Iceland (NE) | ICE_DJU | f | N/A | stranded | 2009 | 14.33 |
| HG569 | Atlantic | Djúpivogur | Iceland (NE) | ICE_DJU | f | N/A | stranded | 2009 | 12.39 |
| HG570 | Atlantic | Djúpivogur | Iceland (NE) | ICE_DJU | f | N/A | stranded | 2009 | 14.31 |
| HG571 | Atlantic | Djúpivogur | Iceland (NE) | ICE_DJU | f | N/A | stranded | 2009 | 14.37 |
| HG573 | Atlantic | Djúpivogur | Iceland (NE) | ICE_DJU | m | N/A | stranded | 2009 | 14.92 |
| HG611 | Atlantic | Froan | Norway | NOR_FRO | f | yearling | live | 1996 | 57.66 |
| HG612 | Atlantic | Froan | Norway | NOR_FRO | m | yearling | live | 1996 | 17.87 |
| HG614 | Atlantic | Froan | Norway | NOR_FRO | m | yearling | live | 1996 | 14.20 |
| HG615 | Atlantic | Froan | Norway | NOR_FRO | f | yearling | live | 1996 | 13.73 |
| HG616 | Atlantic | Froan | Norway | NOR_FRO | m | yearling | live | 1996 | 14.15 |
| HG617 | Atlantic | Froan | Norway | NOR_FRO | f | yearling | live | 1996 | 20.41 |
| HG618 | Atlantic | Froan | Norway | NOR_FRO | m | yearling | live | 1996 | 19.88 |
| HG619 | Atlantic | Froan | Norway | NOR_FRO | m | yearling | live | 1996 | 16.40 |
| HG620 | Atlantic | Froan | Norway | NOR_FRO | f | yearling | live | 1996 | 23.21 |
| HG528 (exluded) | Atlantic | Helgoland | Germany | GER_HEL | m | N/A | live | 2015 | 14.44 |
| HG529 (excluded) | Atlantic | Helgoland | Germany | GER_HEL | m | N/A | live | 2015 | 13.57 |
| HG530 (excluded) | Atlantic | Helgoland | Germany | GER_HEL | m | N/A | live | 2015 | 14.65 |
| HG145 | Atlantic | Isle of May | UK | UK_ILM | f | yearling | live | 2011 | 55.55 |
| HG148 | Atlantic | Isle of May | UK | UK_ILM | f | yearling | live | 2011 | 14.49 |
| HG152 | Atlantic | Isle of May | UK | UK_ILM | m | yearling | live | 2011 | 15.47 |
| HG153 | Atlantic | Isle of May | UK | UK_ILM | f | yearling | live | 2011 | 15.52 |
| HG582 | Atlantic | Isle of May | UK | UK_ILM | f | yearling | live | 2011 | 22.71 |
| HG601 | Atlantic | Kola | Russia | RUS_KOL | m | yearling | live | 1994 | 19.23 |
| HG602 | Atlantic | Kola | Russia | RUS_KOL | m | yearling | live | 1994 | 20.74 |
| HG604 | Atlantic | Kola | Russia | RUS_KOL | m | yearling | live | 1994 | 13.86 |
| HG605 | Atlantic | Kola | Russia | RUS_KOL | f | yearling | live | 1994 | 12.63 |
| HG606 | Atlantic | Kola | Russia | RUS_KOL | m | yearling | live | 1994 | 14.17 |
| HG607 | Atlantic | Kola | Russia | RUS_KOL | f | yearling | live | 1994 | 21.76 |
| HG608 | Atlantic | Kola | Russia | RUS_KOL | m | yearling | live | 1994 | 21.95 |
| HG609 | Atlantic | Kola | Russia | RUS_KOL | m | yearling | live | 1994 | 57.36 |
| HG610 | Atlantic | Kola | Russia | RUS_KOL | m | yearling | live | 1994 | 20.22 |
| HG574 | Atlantic | Patreksfjörður | Iceland (NW) | ICE_PAT | m | N/A | stranded | 2009 | 14.79 |
| HG547 | Atlantic | Strandir | Iceland (NW) | ICE_STR | f | N/A | stranded | 2009 | 15.98 |
| HG551 | Atlantic | Strandir | Iceland (NW) | ICE_STR | m | N/A | stranded | 2009 | 18.97 |
| HG553 | Atlantic | Strandir | Iceland (NW) | ICE_STR | m | N/A | stranded | 2009 | 14.05 |
| HG554 | Atlantic | Strandir | Iceland (NW) | ICE_STR | m | N/A | stranded | 2009 | 44.53 |
| HG703 | Atlantic | Thyborøn | Denmark | DEN_THY | m | juvenile | live | 2022 | 22.24 |
| HG704 | Atlantic | Thyborøn | Denmark | DEN_THY | f | juvenile | live | 2022 | 16.05 |
| HG705 | Atlantic | Thyborøn | Denmark | DEN_THY | m | adult | live | 2022 | 15.48 |
| HG706 | Atlantic | Thyborøn | Denmark | DEN_THY | m | juvenile | live | 2022 | 16.23 |
| HG707 | Atlantic | Thyborøn | Denmark | DEN_THY | m | adult | live | 2022 | 15.31 |
| HG708 | Atlantic | Thyborøn | Denmark | DEN_THY | m | juvenile | live | 2022 | 14.77 |
| HG709 | Atlantic | Thyborøn | Denmark | DEN_THY | m | juvenile | live | 2022 | 18.32 |
| HG710 | Atlantic | Thyborøn | Denmark | DEN_THY | m | juvenile | live | 2022 | 19.05 |
| HG711 | Atlantic | Thyborøn | Denmark | DEN_THY | m | adult | live | 2022 | 16.63 |
| HG712 | Atlantic | Thyborøn | Denmark | DEN_THY | m | adult | live | 2022 | 12.89 |
| HG713 | Atlantic | Thyborøn | Denmark | DEN_THY | f | juvenile | live | 2022 | 16.78 |
| HG714 | Atlantic | Thyborøn | Denmark | DEN_THY | f | yearling | live | 2022 | 15.55 |
| HG715 | Atlantic | Thyborøn | Denmark | DEN_THY | m | adult | live | 2022 | 14.80 |
| HG717 | Atlantic | Thyborøn | Denmark | DEN_THY | f | juvenile | live | 2022 | 14.42 |
| HG718 | Atlantic | Thyborøn | Denmark | DEN_THY | f | yearling | live | 2022 | 36.64 |
| HG720 | Atlantic | Thyborøn | Denmark | DEN_THY | f | adult | live | 2022 | 11.99 |
| HG517 | Atlantic | Thyborøn Rønland Sandø | Denmark | DEN_THY_RON | f | yearling | live | 2015 | 12.27 |
| HG651 | Baltic | Bothnian Bay | Finland | FIN_BOB | m | N/A | hunted | 2017 | 14.17 |
| HG656 | Baltic | Bothnian Bay | Finland | FIN_BOB | f | N/A | hunted | 2017 | 42.30 |
| HG657 | Baltic | Bothnian Bay | Finland | FIN_BOB | m | N/A | hunted | 2017 | 14.50 |
| HG658 | Baltic | Bothnian Bay | Finland | FIN_BOB | f | N/A | hunted | 2017 | 14.36 |
| HG159 | Baltic | Christiansø | Denmark | DEN_CHR | m | adult | live | 2012 | 14.17 |
| HG160 | Baltic | Christiansø | Denmark | DEN_CHR | m | adult | live | 2012 | 14.17 |

| | | | | | | | | | |
|---|---|---|---|---|---|---|---|---|---|
| HG161 | Baltic | Christiansø | Denmark | DEN_CHR | m | adult | live | 2012 | 13.78 |
| HG162 | Baltic | Christiansø | Denmark | DEN_CHR | m | adult | live | 2012 | 14.16 |
| HG163 | Baltic | Christiansø | Denmark | DEN_CHR | m | adult | live | 2012 | 13.85 |
| HG502 | Baltic | Christiansø | Denmark | DEN_CHR | m | juvenile | live | 2016 | 56.53 |
| HG503 | Baltic | Christiansø | Denmark | DEN_CHR | f | juvenile | live | 2016 | 20.83 |
| HG506 | Baltic | Christiansø | Denmark | DEN_CHR | f | juvenile | live | 2016 | 56.52 |
| HG507 | Baltic | Christiansø | Denmark | DEN_CHR | m | juvenile | live | 2016 | 21.97 |
| HG508 | Baltic | Christiansø | Denmark | DEN_CHR | m | juvenile | live | 2016 | 20.22 |
| HG509 | Baltic | Christiansø | Denmark | DEN_CHR | m | juvenile | live | 2016 | 13.13 |
| HG510 | Baltic | Christiansø | Denmark | DEN_CHR | f | juvenile | live | 2016 | 16.76 |
| HG511 | Baltic | Christiansø | Denmark | DEN_CHR | m | juvenile | live | 2016 | 15.84 |
| HG521 | Baltic | Christiansø | Denmark | DEN_CHR | f | juvenile | live | 2014 | 18.49 |
| HG522 | Baltic | Christiansø | Denmark | DEN_CHR | m | juvenile | live | 2014 | 17.51 |
| HG523 | Baltic | Christiansø | Denmark | DEN_CHR | f | juvenile | live | 2014 | 15.81 |
| HG524 | Baltic | Christiansø | Denmark | DEN_CHR | f | N/A | live | 2016 | 20.90 |
| HG525 | Baltic | Christiansø | Denmark | DEN_CHR | m | N/A | live | 2016 | 15.98 |
| HG526 | Baltic | Christiansø | Denmark | DEN_CHR | f | N/A | live | 2016 | 17.93 |
| HG527 | Baltic | Christiansø | Denmark | DEN_CHR | m | N/A | live | 2016 | 15.96 |
| HG167 | Baltic | Falsterbo | Sweden | SWE_FAL | m | juvenile | live | 2012 | 14.85 |
| HG168 | Baltic | Falsterbo | Sweden | SWE_FAL | m | juvenile | live | 2012 | 14.86 |
| HG169 | Baltic | Falsterbo | Sweden | SWE_FAL | f | juvenile | live | 2012 | 14.57 |
| HG170 | Baltic | Falsterbo | Sweden | SWE_FAL | m | juvenile | live | 2012 | 14.92 |
| HG171 | Baltic | Falsterbo | Sweden | SWE_FAL | m | juvenile | live | 2012 | 14.89 |
| HG652 | Baltic | Gulf of Finland | Finland | FIN_GOF | m | N/A | hunted | 2017 | 13.25 |
| HG653 (exluded) | Baltic | Gulf of Finland | Finland | FIN_GOF | f | N/A | hunted | 2017 | 14.44 |
| HG654 | Baltic | Gulf of Finland | Finland | FIN_GOF | f | N/A | hunted | 2017 | 13.45 |
| HG655 | Baltic | Gulf of Finland | Finland | FIN_GOF | f | N/A | hunted | 2017 | 14.11 |
| HG659 | Baltic | Gulf of Finland | Finland | FIN_GOF | f | N/A | hunted | 2017 | 14.35 |
| HG641 | Baltic | Gulf of Riga | Estonia | EST_GOR | m | pup | live | 2016 | 43.82 |
| HG642 | Baltic | Gulf of Riga | Estonia | EST_GOR | f | pup | live | 2016 | 14.18 |
| HG643 | Baltic | Gulf of Riga | Estonia | EST_GOR | f | pup | live | 2016 | 14.09 |
| HG644 | Baltic | Gulf of Riga | Estonia | EST_GOR | f | pup | live | 2016 | 12.99 |
| HG645 | Baltic | Gulf of Riga | Estonia | EST_GOR | f | pup | live | 2016 | 14.31 |
| HG007 | Baltic | Rødsand | Denmark | DEN_ROD | m | N/A | stranded | 2000 | 14.96 |
| HG008 | Baltic | Rødsand | Denmark | DEN_ROD | m | N/A | stranded | 2009 | 14.49 |
| HG009 | Baltic | Rødsand | Denmark | DEN_ROD | m | N/A | stranded | 2001 | 14.70 |
| HG155 | Baltic | Rødsand | Denmark | DEN_ROD | f | juvenile | live | 2009 | 10.16 |
| HG300 | Baltic | Rødsand | Denmark | DEN_ROD | m | pup | live | 2013 | 14.36 |
| HG301 | Baltic | Rødsand | Denmark | DEN_ROD | m | pup | live | 2013 | 42.87 |
| HG302 | Baltic | Rødsand | Denmark | DEN_ROD | f | pup | live | 2013 | 13.96 |
| HG303 | Baltic | Rødsand | Denmark | DEN_ROD | f | pup | live | 2019 | 14.20 |
| HG304 | Baltic | Rødsand | Denmark | DEN_ROD | f | pup | live | 2019 | 12.53 |
| HG305 | Baltic | Rødsand | Denmark | DEN_ROD | m | pup | live | 2019 | 14.19 |
| HG306 | Baltic | Rødsand | Denmark | DEN_ROD | m | pup | live | 2024 | 14.28 |
| HG307 | Baltic | Rødsand | Denmark | DEN_ROD | f | pup | live | 2024 | 14.03 |
| HG308 | Baltic | Rødsand | Denmark | DEN_ROD | f | pup | live | 2024 | 13.99 |
| HG309 | Baltic | Rødsand | Denmark | DEN_ROD | m | pup | live | 2024 | 26.18 |
| HG310 | Baltic | Rødsand | Denmark | DEN_ROD | f | pup | live | 2024 | 13.94 |
| HG311 | Baltic | Rødsand | Denmark | DEN_ROD | m | pup | live | 2024 | 14.68 |
| HG513 | Baltic | Rødsand | Denmark | DEN_ROD | m | pup | live | 2014 | 10.43 |
| HG514 | Baltic | Rødsand | Denmark | DEN_ROD | f | pup | live | 2014 | 14.48 |
| HG515 | Baltic | Rødsand | Denmark | DEN_ROD | f | pup | live | 2014 | 13.94 |
| HG516 | Baltic | Rødsand | Denmark | DEN_ROD | m | pup | live | 2014 | 15.18 |
| HG625 | Baltic | Stockholm Archipelago | Sweden | SWE_STA | f | N/A | hunted | 2010 | 12.60 |
| HG626 | Baltic | Stockholm Archipelago | Sweden | SWE_STA | f | N/A | hunted | 2010 | 14.23 |
| HG627 | Baltic | Stockholm Archipelago | Sweden | SWE_STA | m | N/A | hunted | 2010 | 12.82 |
| HG628 | Baltic | Stockholm Archipelago | Sweden | SWE_STA | m | N/A | hunted | 2011 | 13.90 |
| HG629 | Baltic | Stockholm Archipelago | Sweden | SWE_STA | m | N/A | hunted | 2011 | 14.26 |
